# Directed brain connectivity identifies widespread functional network changes in Parkinson’s disease

**DOI:** 10.1101/2021.01.04.425206

**Authors:** Mite Mijalkov, Giovanni Volpe, Joana B. Pereira

## Abstract

Parkinson’s disease (PD) is a progressive neurodegenerative disorder characterized by topological changes in large-scale functional brain networks. These networks are commonly analysed using undirected correlations between the activation signals of brain regions. However, this approach suffers from an important drawback: it assumes that brain regions get activated at the same time, despite previous evidence showing that brain activation features causality, with signals being typically generated in one region and then propagated to other ones. Thus, in order to address this limitation, in this study we developed a new method to assess whole-brain directed functional connectivity in patients with PD and healthy controls using anti-symmetric delayed correlations, which capture better this underlying causality. To test the potential of this new method, we compared it to standard connectivity analyses based on undirected correlations. Our results show that whole-brain directed connectivity identifies widespread changes in the functional networks of PD patients compared to controls, in contrast to undirected methods. These changes are characterized by increased global efficiency, clustering and transitivity as well as lower modularity. In addition, changes in the directed connectivity patterns in the precuneus, thalamus and superior frontal gyrus were associated with motor, executive and memory deficits in PD patients. Altogether, these findings suggest that directional brain connectivity is more sensitive to functional network changes occurring in PD compared to standard methods. This opens new opportunities for the analysis of brain connectivity and the development of new brain connectivity markers to track PD progression.

## I. INTRODUCTION

Parkinson’s disease (PD) is a complex neurodegenerative disorder characterized by a wide range of motor and non-motor symptoms such as memory, executive, visuospatial or olfactory deficits [13, 32]. The presence of such diverging symptoms suggests that the brain changes occurring in PD cannot be directly linked to the dysfunction of a single brain region but rather to widespread changes in functional connectivity between many regions or brain networks [47].

Functional connectivity can be measured using functional magnetic resonance imaging (MRI), a non-invasive technique that detects changes in blood oxygen level dependent (BOLD) signals, which are considered to reflect the underlying neuronal brain activity [7]. In PD patients, several studies have shown that motor and non-motor symptoms can arise due to the loss of integrity in these functional connections [21, 58]. In particular, abnormal functional connectivity in the basal ganglia–thalamocortical network [4, 9, 27] has been linked to motor symptoms in PD, whereas changes in the default mode, dorsal-attention, fronto-parietal, salience and associative visual networks [1, 3, 21, 25, 49, 59, 63] have been shown to correlate with cognitive and executive deficits in these patients.

In the past few years, several studies have used functional MRI to assess the functional brain connectome, a whole-brain network that summarizes the complete set of pairwise functional connections in the brain [8]. This network consists of set of nodes, or brain regions, connected by edges, representing the strength of the functional connections. This connectivity network can then be analyzed using graph theory by computing several global and local measures that reflect whether brain regions are efficiently connected by short network paths (global efficiency) or are well integrated into their neighborhood (clustering) or community (modularity) [53]. These analyses have shown significant changes in the global efficiency, local efficiency and clustering coefficient in the whole brain [2, 26] or within specific networks in PD patients [36, 39, 64]. Changes in the nodal network topology of prefrontal and supplementary motor areas as well as the striatum and thalamus [16, 38, 55, 67] have also been reported in PD patients, sometimes in association with clinical measures [37, 56].

Despite being useful to assess network changes in PD, these studies were based on the assumption that brain activity in different brain regions occurs simultaneously and therefore can be captured by same-time undirected correlations in the activation signals between them. As such, they do not convey information about the directionality of the interaction between brain regions [20], which is important due to an increasing number of studies showing that directed brain activity patterns are altered in PD. These directed patterns have been assessed using dynamic causal modelling [33, 52], structural equation modelling [45, 51], psycho-physiological interactions [68] or Granger causality [22, 66] methods. Due to the complex nature and longer computational time required by these methods, their application is currently limited to the assessment of brain connectivity between a few regions or to the analysis of functional MRI data acquired during a specific task, which normally relies on a priori hypotheses of which brain regions should be tested. Moreover, several generalizations for the assessment of directed whole-brain connectivity have also been recently proposed [18, 19, 24, 48, 50], however their applicability to neurodegenerative diseases has not been evaluated and, to our knowledge, they do not provide connectivity information from multiple timescales. Therefore, an intuitive and computationally light method that can analyze whole-brain directed resting-state connectivity patterns at different timescales is currently missing.

Here, we present a method based on anti-symmetric lagged correlations to assess restingstate, whole-brain directed functional networks. First, we obtain a lagged correlation adjacency matrix for each patient by calculating the pairwise lagged correlations between all pairs of brain regions. Then, the anti-symmetric correlations was derived as the anti-symmetric part of the lagged correlation adjacency matrix. We demonstrate that the topological organization of these functional networks is more sensitive to pathological changes related to PD when compared to functional networks built by standard undirected methods.

## II. RESULTS

### A. Construction of directed functional networks

Due to previous evidence showing temporal lags in the activation signals between connected brain regions [23, 31, 43], we calculated directed functional connectivity between brain regions using lagged Pearson’s correlations. In this approach, a brain region is considered to have a directed interaction with other brain regions if its activation time series has similar properties with the time-shifted version of the second brain region’s activation pattern. Moreover, as brain activity is a dynamic process that changes over time [29], we evaluate directed functional connectivity at multiple temporal lags, thereby exploring the functional activation of the brain at different time scales *(“Methods: Lagged correlation”)*.

Figure 1 illustrates the different methods we use to calculate the functional connectivity networks for a set of 5 brain regions and their activation time series (Fig. 1a). The connectivity matrix and the corresponding network calculated by the lagged correlation adjacency method for these 5 brain regions are shown in Fig. 1b. It shows that the lagged correlation method evaluates the directed connection between two regions in both directions; a pair of elements in the lagged adjacency matrix (namely, (*i, j*) and (*j, i*)) encode the estimated directed influence of brain region *i* to brain region *j* and vice versa. As any other square matrix, the lagged correlation adjacency matrix can be uniquely expressed as the sum of a symmetric and anti-symmetric matrix. Specifically, the anti-symmetric matrix captures the directionality of the functional network, identifying the relevant directed connections between the couples of brain regions (Fig. 1c). We call this method “anti-symmetric correlation” *(“Methods: Anti-symmetric correlation”)*.

**FIG. 1.**
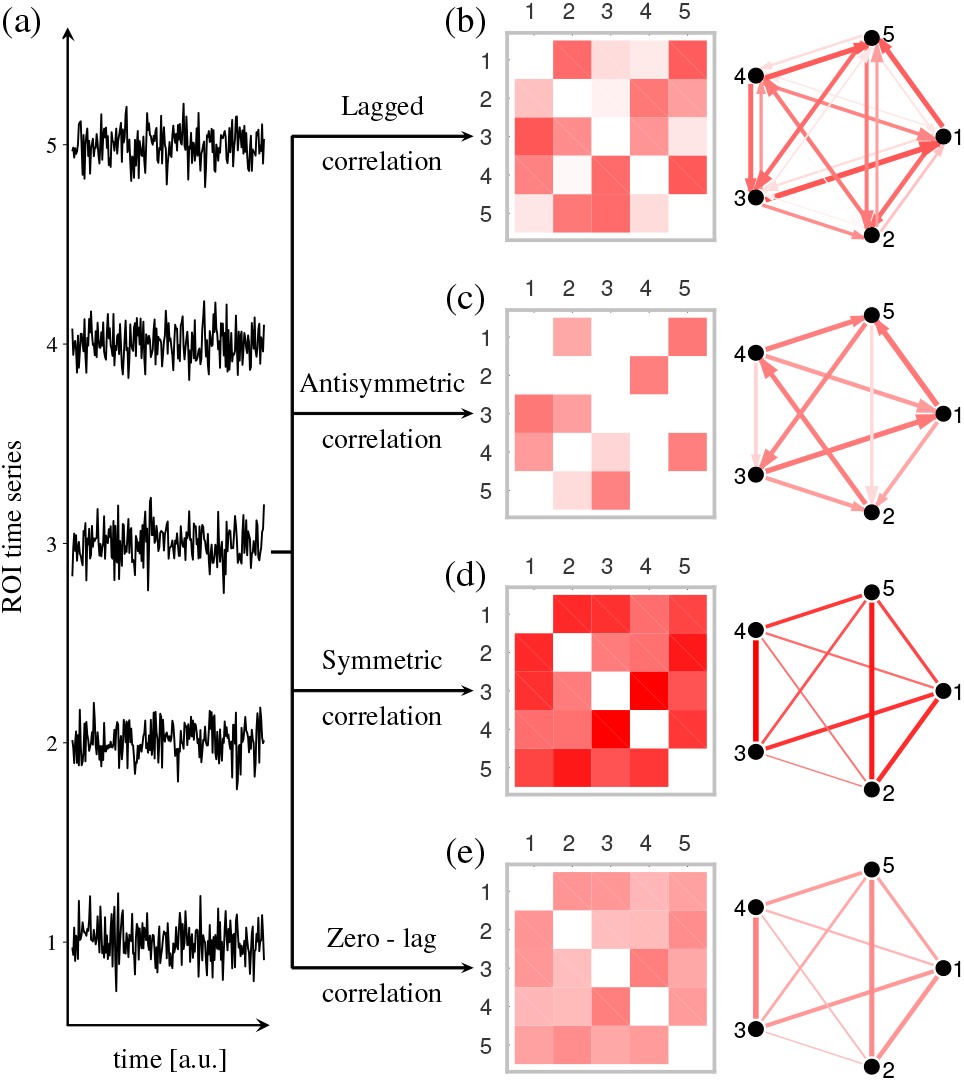
Different methods used to calculate functional networks. (a) For illustration purposes, we show an example of the time activation series of only 5 nodes. (b) Lagged correlation functional networks can be estimated by calculating the lagged Pearson’s correlation coefficient between these time series, at different lags. Here, the lagged adjacency matrix and corresponding network are calculated at lag of 1. The lagged adjacency matrix can be written as a sum of a (c) anti-symmetric and (d) symmetric matrices. Finally, for comparison we show the commonly used method of zero-lag correlation (e). In all matrices, redder colors and thicker lines indicate stronger connections.

To highlight the effectiveness of the directed networks in detecting topological changes between controls and PD patients, we compare our method with two undirected network approaches. In the first approach, functional connectivity is evaluated as the symmetric matrix extracted from the lagged correlation adjacency matrix (Fig. 1d), in which the undirected connection between two regions is the sum of the weights of the two corresponding directed connections *(“Methods: Symmetric correlation”)*. Secondly, we also compare our method with the conventional approach to quantify functional connectivity, in which the connectivity strength between two regions is estimated by calculating the zero-lag Pearson’s correlation coefficient (Fig. 1e) between their activation time series *(“Methods: Zero-lag correlation”)*.

We tested the ability of all 4 methods to detect topological changes in PD patients in a cohort of 95 PD patients and 15 controls with functional MRI scans from the Parkinson’s Progression Markers Initiative *(“Methods: Participants”)*. The nodes in the adjacency matrices corresponded to the 200 brain regions derived from the Craddock atlas [14], while the edges were calculated according to the four methods described above, yielding 4 different weighted adjacency matrix for each participant. For each adjacency matrix, we calculated a binary matrix where the correlation coefficient was considered 1 if it was above a certain threshold, and 0 if it was below. We performed the thresholding at the complete available range of network densities (D) of the anti-symmetric correlation (*D*_*min*_ = 1%, to *D*_*max*_ = 50% in steps of 1%) and we compared the network topologies across that range. The negative correlation coefficients and self-connections were excluded from the analysis by setting them to zero.

### B. Average group networks show different behavior across different temporal lags

We calculated group-representative adjacency matrices at different temporal lags by averaging the weighted, subject-specific adjacency matrices. The histograms of the connection weights are shown in Figure 2 for the lagged (Fig. 2a), anti-symmetric (Fig. 2b) and symmetric (Fig. 2c) correlations as a function of different temporal lags. Fig. 2 shows a general decrease of the strength of directed connectivity in PD patients for all analyses at all lags when compared to healthy controls. Furthermore, in PD patients, we observe that the connectivity strength distribution becomes narrower with increasing temporal lags for all analyses. This observation indicates that, with higher temporal lags, more nodes have similar functional connectivity strength. Therefore, large temporal lags are unsuitable for the analysis of between-group topological differences because they cannot capture any variations in the directional flow in the network, restricting our analysis to small temporal lags in the range 1 − 7.

**FIG. 2.**
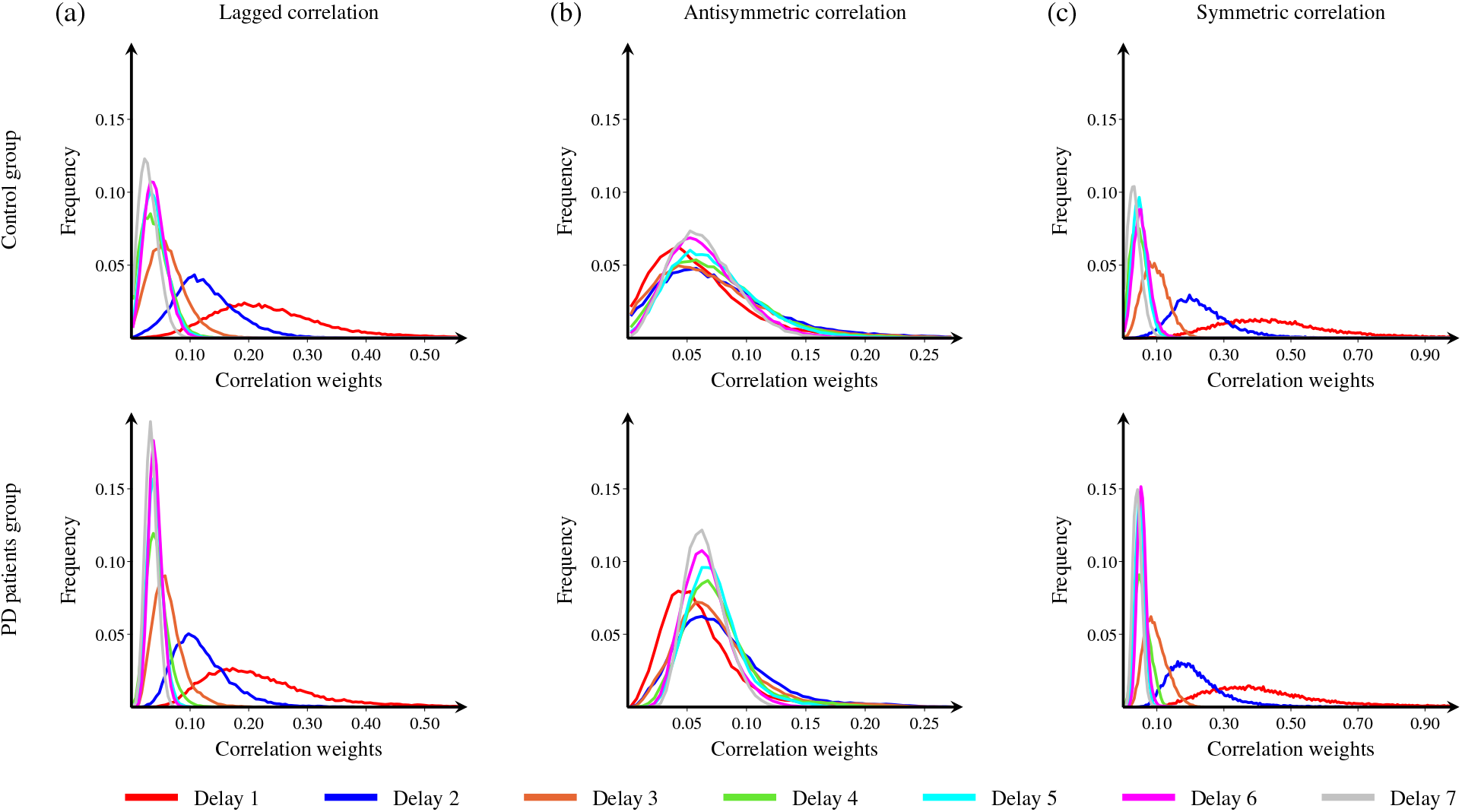
Connectivity strength distribution at different temporal lags. Histograms show the distribution of connectivity strengths of the average adjacency matrices for controls (top row) and PD patients (bottom row) as a function of different temporal lags. The individual subject connectivity matrices were calculated using (a) lagged analyses, (b) anti-symmetric analyses and (c) symmetric analyses. Only the lags used in the analysis are shown in this figure.

### C. Differences in global topology

To assess the ability of these methods to detect global network changes between patients and controls, we calculated the global efficiency, local efficiency, clustering coefficient, transitivity and modularity (Fig. 3, left to right columns). Only the anti-symmetric correlation method showed widespread significant differences between PD patients and controls in network measures; this entails that the differences are contained in the anti-symmetric part of the lagged correlation matrix. These differences consisted of increases in the clustering coefficient and transitivity in the PD group compared with controls at several higher network densities (clustering coefficient: 16% − 50%; transitivity: 20% − 50%). The global and local efficiency also showed differences between patients and controls, being increased in PD patients across most network densities (global efficiency: 2% − 50%; local efficiency: 6% − 50%). Finally, we also found significant decreases in the modularity in PD patients compared with controls, which were only present at higher network densities (21% − 50%). In contrast, the undirected and lagged correlation methods did not show any significant differences between any of the global network measures between PD patients and controls. Fig. 3 summarizes the results obtained for the temporal lag 1; the corresponding results for lags 2 − 7 are shown in supplementary figures S1 - S6.

**FIG. 3.**
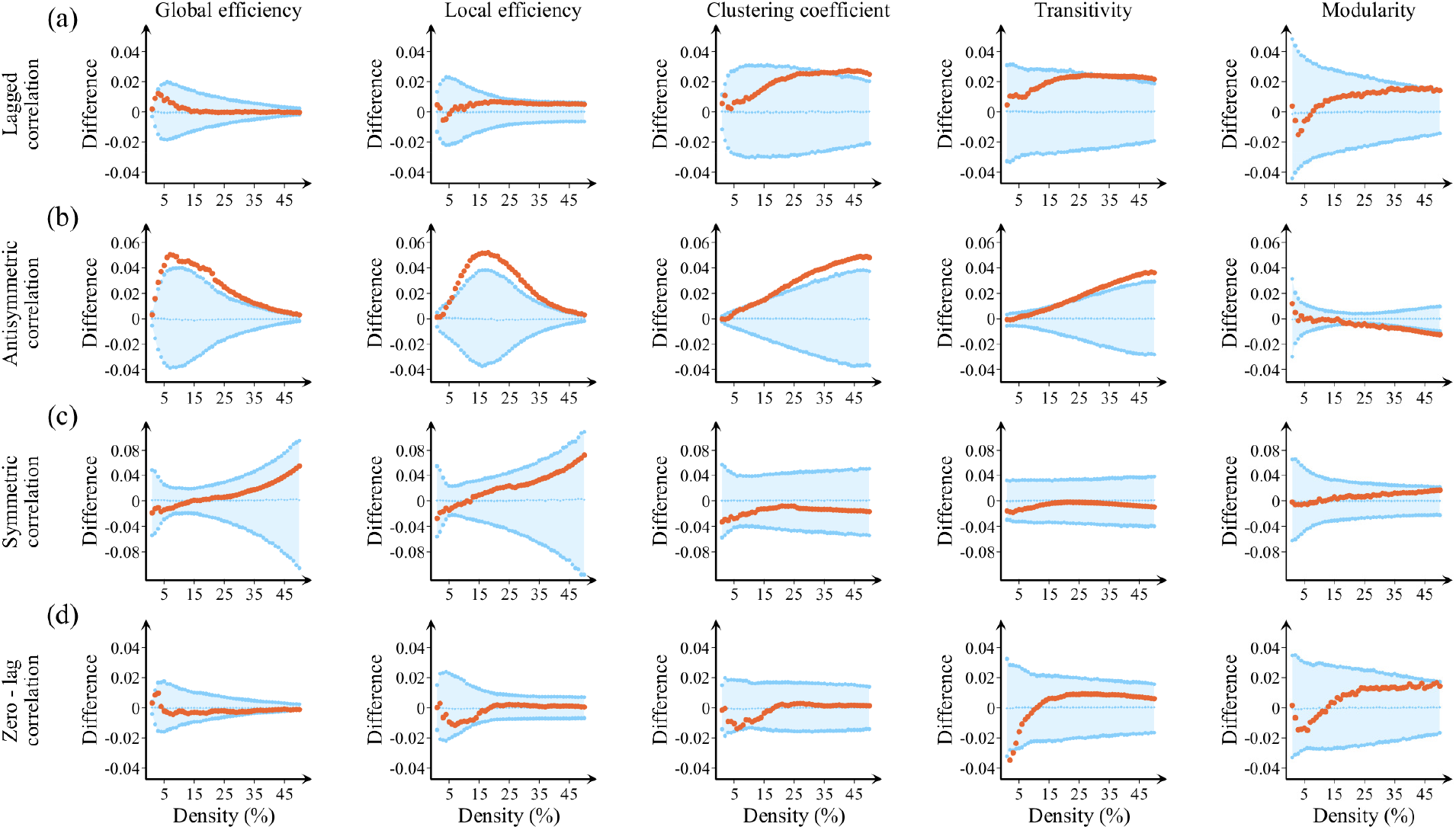
Differences between controls and PD patients in global network measures. Plots showing the differences between controls and PD patients in the global efficiency, local efficiency, clustering coefficient, transitivity and modularity using (a) lagged correlation, (b) anti-symmetric correlation, (c) symmetric correlation and (d) zero-lag correlation methods. The plots show the upper and lower bounds of the 95% confidence intervals (CI) in blue, and the differences in the network measures between groups in orange circles as a function of network density. The differences are considered statistically significant if they fall outside the CIs.

### D. Differences in nodal topology

Furthermore, using the anti-symmetric correlation method, we also identified directed connectivity changes in several brain regions in PD patients compared to controls (Fig. 4). These regions included the precuneus, which showed increases in the in-global efficiency and decreases of the out-degree at various temporal lags; the thalamus, which showed decreases of overall connectivity in PD patients at lags 4 and 5; the superior frontal gyrus, which showed an increased outflow connectivity in PD patients; and the fusiform, which showed increases in the in-global efficiency in PD patients at lag 7. The other three analysis methods were not able to identify any significant between-group differences.

**FIG. 4.**
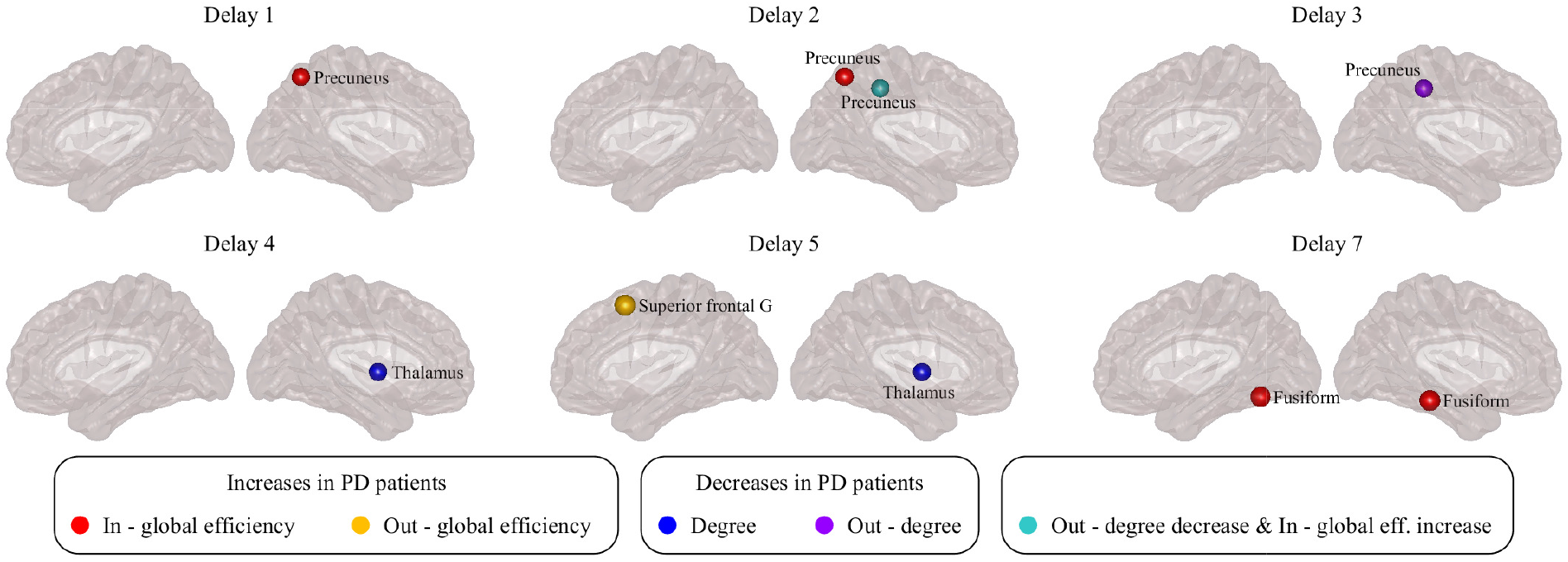
Differences between controls and PD patients in nodal network measures. Visual display of the nodes that show significant differences between controls and PD patients in network measures using the anti-symmetric correlation method. For simplicity, here we show only the regions that were significant at the middle network density *D* = 25% at the temporal lags indicated in the figure. Differences between groups were evaluated using non-parametric permutation tests. Only regions that show significant differences after correcting for multiple comparisons (FDR at q=0.05) are plotted.

### E. Correlation analysis with clinical measures in PD patients

All global network measures were significantly associated with the UPDRS-III motor scores and executive scores (Letter-Number sequencing test) across all lags. In addition, the clustering and transitivity also correlated with executive scores (symbol digit modalities test) at lag 1, whereas global efficiency correlated with memory (Hopkins verbal learning test) at lag 5. Global and local efficiency, clustering and transitivity correlated with visuospatial scores (Benton’s judgment of line orientation test) at lag 7. The correlation coefficients between the global measures and corresponding clinical test scores and the associated p-values that remained significant after adjusting for multiple comparisons (FDR, *q* = 0.05) are summarized in Supplementary Tables S1-S17.

Regarding the nodal network measures, the out-degree of the precuneus was significantly associated with olfactory function (UPSIT smell identification test) at lags 2 (p − value = 0.003; r = −0.31) and 3 (p − value = 0.006; r = −0.28). Furthermore, memory (Hopkins verbal learning test) significantly correlated with the out-degree in the precuneus at lag 3 (p − value = 0.048; r = 0.21) as well as the thalamus at lag 4 (p − value = 0.025; r = 0.23). The degree of the thalamus correlated with visuospatial scores (Benton’s judgment of line orientation test) at lags 4 (p − value *<* 0.001; r = −0.36) and 5 (p − value = 0.031; r = −0.23). Finally, the fusiform’s in-global efficiency was associated with the semantic fluency tests at lag 7 (p − value = 0.049; r = 0.21).

### F. Effect of dopaminergic medication on functional network topology

To evaluate the effect of levodopa-equivalent doses on functional network organization we compared the networks of medicated patients to those who were not receiving medication (details about the two subgroups are shown in Table S18). We did not find any differences in the global network topology between these groups. Regarding nodal topology, there were significant increases in the out-global efficiency in the precuneus, superior parietal lobule and superior occipital gyrus at lag 1, significant decreases of the in-degree in the precuneus at lag 2, and increases of the in-degree of thalamus at lags 2 and 3 in medicated patients versus non-medicated ones (Fig. 5). Of note, none of these results overlapped with the results of the main analyses.

**FIG. 5.**
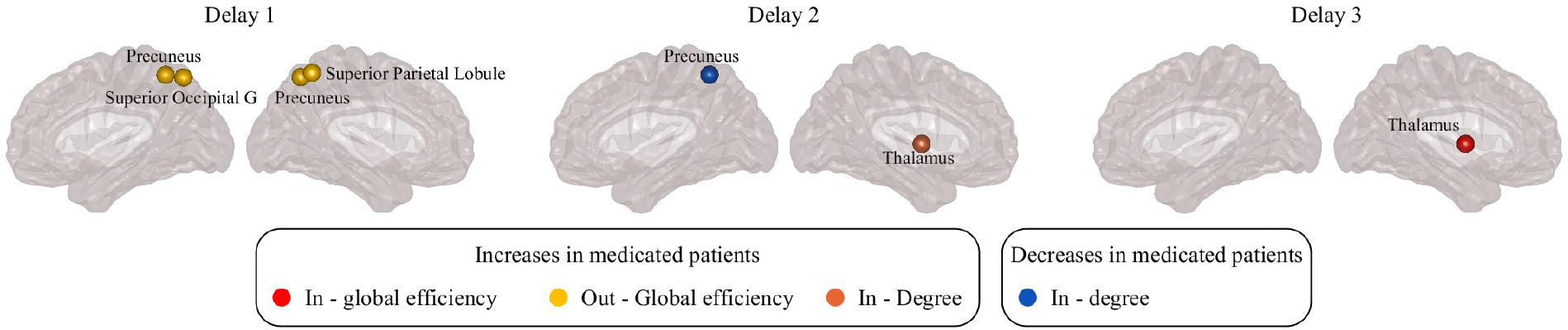
Differences between medicated and non-medicated PD patients in nodal network measures. Plots showing the differences between medicated and non-medicated PD patients in nodal network measures evaluated at network density *D* = 25% at different temporal lags, in the case of anti-symmetric correlation. Differences between groups were evaluated using non-parametric permutation test. Only regions that show significant differences after correcting for multiple comparisons (FDR at q=0.05) are plotted.

### G. Influence of mild cognitive impairment on functional network topology

Due to previous evidence showing that PD patients with mild cognitive impairment (MCI) show more widespread network changes compared to cognitively normal patients [2, 25], we performed an additional analysis to compare these two groups (patient characteristics for both subgroups are shown in Table S19). Only one significant difference was found in the cerebellum, which showed significant degree decreases in patients with MCI at lag 3 compared to cognitively normal patients.

## III. DISCUSSION

In this study we propose a new method to analyze directed functional connectivity that uses the information stored in the temporal lags between the activation of brain regions. To our knowledge, there are currently no methods that allow assessing directed functional connectivity across the entire brain at multiple timescales and studying the corresponding topological changes. Our anti-symmetric correlation method was developed to address this gap, showing that whole-brain directed connectivity is useful to characterize the connectomes of patients with PD by detecting widespread functional alterations that were not identifiable by conventional zero-lag methods. In addition, we found that the changes identified by the anti-symmetric correlation method were associated with motor, executive and memory deficits in patients, suggesting they are clinically meaningful. Altogether, our findings indicate that the directional flow in brain activation signals contains exclusive information that is not captured by other methods, and could potentially be used as a new marker of functional network changes in PD.

Functional connectivity describes the statistical dependencies in the activation patterns between brain regions, and is closely associated with behavior and cognitive functions [62]. Such statistical dependencies can be quantified using a vast amount of measures derived from graph theory, which typically consider two regions to be connected if the Pearson correlation between their activation signals is strong. However, this method is hindered by the fact that it only captures linear, simultaneous and undirected dependencies between brain regions. There is evidence showing that the relationship between brain regions is not always linear and that there are often delays between their activation signals [23, 31, 43]. Thus, capturing the information stored in these temporal delays or lags is crucial to obtain a more accurate characterization of the brain’s functional connectivity.

To demonstrate that the anti-symmetric correlation method is useful to characterize functional connectivity, we tested its performance on a cohort of PD patients and healthy controls. Our method detected an abnormal global topology in the functional connectomes of PD patients, characterized by increases in global efficiency, local efficiency, clustering and transitivity as well as decreases in the modularity when compared to healthy controls. The increases in global efficiency can be interpreted in light of previous studies showing that brain networks with a random organization have shorter network paths and greater global efficiency [57]. In addition to PD patients [16, 38, 61], this phenomenon has been shown to also occur in the networks of patients with schizophrenia [17] and Alzheimer’s disease [60], being associated with executive impairment and other cognitive deficits [57]. On the other hand, the increases of clustering and transitivity in the networks of PD patients indicate an increase in the number of directed cyclic connections within local neighborhoods. This formation of closed triangles between neighboring regions increases the segregation and fragmentation of the functional networks, which has also been reported in some studies in patients with PD [2, 26]. These changes were accompanied by lower modularity, suggesting that the fragmentation occurring in the networks of PD patients did not result in well-defined communities, which is normally regarded as a sign of brain pathology [41]. Thus, our findings show that the changes occurring in PD patients reflect both increased integration and segregation in the directed functional networks. These changes were associated with worse performance on various clinical and cognitive tests measuring motor function, executive abilities, memory and visuospatial functions, suggesting that changes in global directed activation patterns can be an indicator of worse clinical progression in PD.

In addition to global network changes, we also observed alterations in the topology of specific brain regions. For instance, the precuneus showed an increase in the in-global efficiency and a decrease in the out-degree, which were associated with memory and olfactory deficits. These findings are in line with previous evidence showing that the precuneus is a brain hub that plays an important role in memory, attention and other cognitive functions [12]. Several studies have shown changes in the functional connectivity patterns of the precuneus in PD patients [15, 16, 26, 63]. Our findings offer an additional insight into the nature of these alterations. In particular, they indicate a specific shift to an increased number of in-coming connections accompanied by a decrease in the number of outgoing connections. This imbalance between in- and out-connectivity could possibly alter the role of the precuneus in the patients’ networks, making it an inefficient hub. Furthermore, this abnormal local topology could result in changes in the connectivity patterns within DMN and its strong connections with the olfactory system [34], leading to deficits in memory and loss of smell commonly experienced by PD patients.

Moreover, an increase of the in-global efficiency was observed in the fusiform gyrus, which was associated with semantic fluency. Similar changes have been previously observed in the connectivity of the fusiform gyrus, which could lead to deficits in visual processing functions and decreased performance in the verbal fluency tasks [6, 11].

Finally, in our study, the superior frontal gyrus also had an increased outflow connectivity in PD patients, while the thalamus displayed a decreased overall connectivity, which correlated with memory and visuospatial deficits. Such changes in the functional activity of the frontal cortex have been associated with deficits in executive functions in PD patients, for example working memory, cognitive flexibility and problem solving. Due to its strong connections with the striatum, these deficits have also been linked with dysfunction of the frontostriatal networks [44, 46]. Being a part of the basal ganglia thalamo-cortical network, the thalamus carries information from the basal ganglia to the cerebral cortex, making it an important hub in functional brain networks [30]. As such, the thalamus plays an important role in many functions, such as motor abilities, visually-guided actions, learning and memory [54, 65]. Thus, our results provide further support the role of the thalamus in contributing to functional abnormalities in the networks of PD patients and the various motor and non-motor deficits they present.

There is ample evidence showing that brain connectivity is a dynamic process that changes over time [29, 35]. An advantage of the anti-symmetric correlation method is its ability to calculate functional connectomes at different temporal lags, allowing to analyse functional connectivity as a dynamic process that can change across multiple temporal scales. Although we found an uniform global topology across all lags, the changes in regional topology varied substantially between different temporal scales. These results suggest that, in PD patients, the general efficiency in information transfer is maintained at multiple temporal scales by conserving the global topological properties of the functional network. However, as different sets of brain regions co-activate at different temporal lags, the local topology of the regions varies with the value of the lags. As a result, abnormal regional changes are shown in distinct regions at different temporal lags in PD patients compared to controls. The connectivity values of these different regions were associated with worse performance in motor and cognitive tests, suggesting that motor and cognitive deficits in PD patients may be associated with brain connectivity changes occurring at different temporal scales.

In order to assess which temporal scales were most relevant for our analysis, we plotted the connectivity weight profiles of the average connectivity matrices for both controls and PD patients. For large delays, the connection weight histograms of both groups were narrow. This shows that large temporal lags are unable to capture variations in the directed activation flow in the network, instead assigning similar weights to a large number of connections. Therefore, in order to be able to capture this functional variation, we restricted our analysis to small temporal lags in the range 1-7, in agreement with previous studies using multivariate models to analyse directed functional connectivity [22].

Although the current study has several strengths, some limitations should also be recognised, which present opportunities for future work. Although we tested the anti-symmetric correlations on a well characterized sample of PD patients, our findings should be replicated in larger and independent cohorts. In particular, several patients in the current cohort underwent functional MRI while on medication, which has previously been shown to influence brain connectivity [15, 45, 67]. In this study, we assessed the effects of medication on our results by performing correlation analyses between the levodopa-equivalent doses and the topological graph measures, as well as comparing the networks of medicated and non-medicated patient groups. Our analyses showed that there was no association between medication doses and topological measures, and there were no differences in the global measures between the medicated and non-medicated patients. The only significant results that were observed in medicated compared to non-medicated groups was an increase of the out-global efficiency in the superior occipital gyrus, superior parietal lobule and precuneus at lag of 1, a decrease in the in-degree of precuneus at lag 2 and increases in the in-global efficiency and the in-degree of thalamus at lags 2 and 3. Since these regions did not overlap with the measures or regions that showed differences between PD patients and controls in our main analysis, most likely they did not influence our results. In addition, it has also been demonstrated that PD patients with mild cognitive impairment have a different functional connectivity pattern when compared to cognitively normal patients [1–3, 38]. As 20% of the PD patients in our study were diagnosed with MCI, we also performed an additional analysis to compare them with the cognitively normal PD patients. We found no topological differences in the directed functional connectomes between the two groups, suggesting that the presence of MCI also did not affect the main results. This result is in contrast with previous studies showing that the presence of MCI has an effect on network topology in patients with PD [2, 38]. This discrepancy is probably associated with the differences in clinical characteristics between our sample and the cohorts used in previous studies.

Despite these limitations, in this study we show that the information stored in the temporal activation lags can be used to assess the directed connections between all the brain regions of the functional connectome. Our findings show that these directed connections can detect specific topological changes in PD patients at multiple temporal scales, offering increased sensitivity to PD-related changes compared to undirected methods. These findings suggest that our method could potentially be used to improve the diagnosis of PD or identify patients with worse disease progression.

## IV. METHODS

### A. Participants

This study included 95 PD patients and 15 controls from the Parkinson’s Progression Markers Initiative (PPMI) database [40] (Table 1). For up-to-date information on the study, visit www.ppmi-info.org. PD patients were diagnosed within 2 years of the screening visit, were entirely untreated at enrollment, had a Hoehn and Yahr [28] stage of I or II, and were required to have a dopamine transporter deficit on DaTSCAN imaging. Only subjects with a functional MRI scan that passed quality control before and after image preprocessing were included. Motor symptoms were assessed using the unified Parkinson’s disease rating scale (UPDRS) and olfactory function was evaluated using the smell identification test (UPSIT). In addition, all subjects completed several cognitive tests that assessed visuospatial functions (15-item version of the Benton’s judgment of line orientation test), verbal memory (total immediate recall and delayed recall of the Hopkins verbal learning test-revised, HVLT-R), executive functions (the letter number sequencing test, semantic and phonemic fluency tests) and attention (symbol digit modalities test (SDMT). The total levodopa-equivalent doses were recorded for all PD patients.

**TABLE 1.**
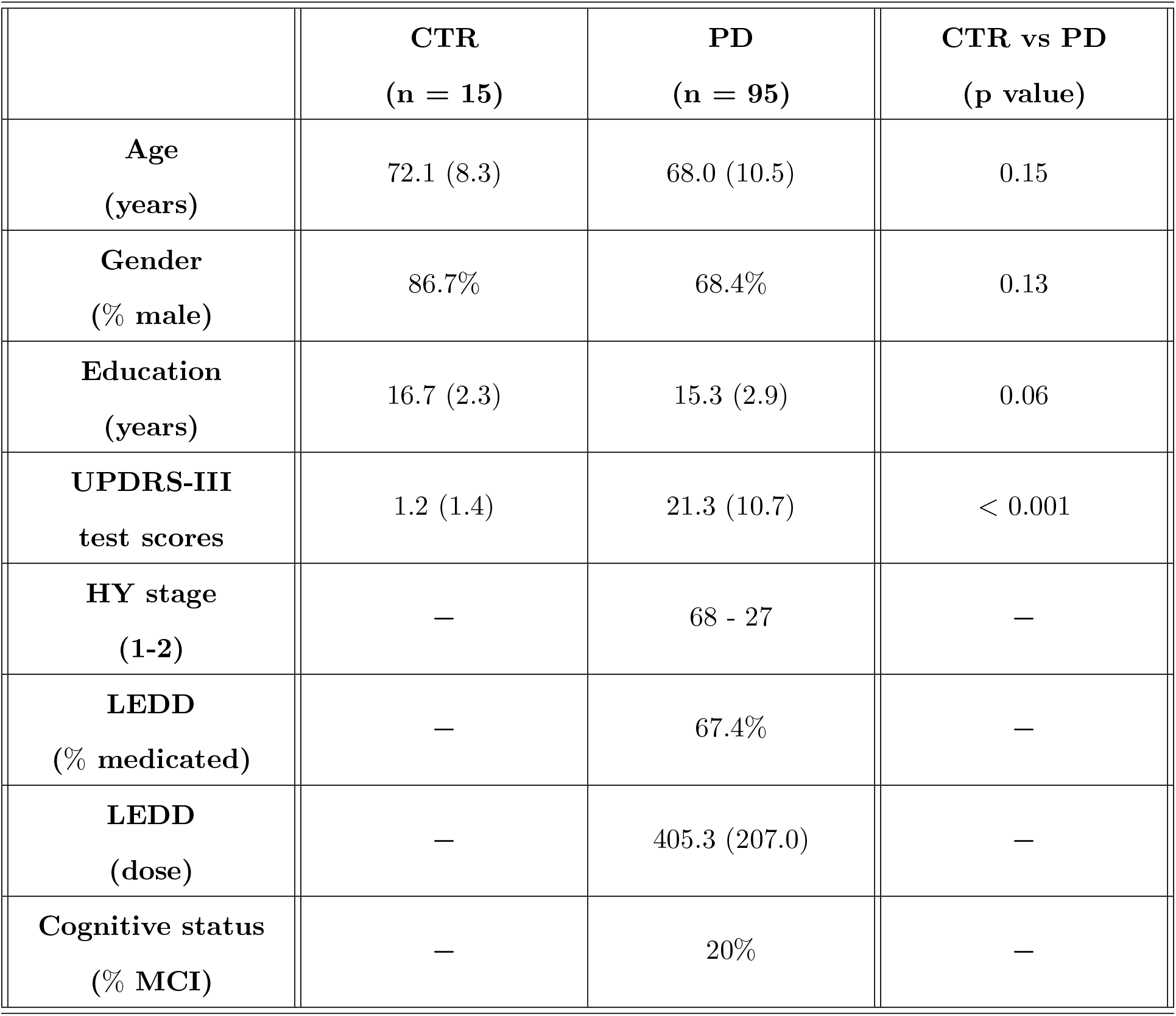
Characteristics of the sample. Means are followed by standard deviation in parenthesis. Permutation tests with 10000 permutations were used to compare groups for age, gender, education, UPDRS-III scores, LEDD dose and Hoehn and Yahr stage. CTR, controls; PD, Parkinson’s disease; UPDRS-III, Unified Parkinson’s disease rating scale–Part III; HY stage, Hoehn and Yahr stage; LEDD, levodopa equivalent dose.

### B. Image acquisition

All subjects were scanned on a 3 Tesla Siemens scanner using an echo planar functional MRI sequence with the following parameters: 212 time points, repetition time = 2400 milliseconds, echo time = 25 milliseconds, field of view = 222 mm, flip angle = 80 degrees and 3.3 mm isotropic voxels. During the scanning session, subjects were instructed to keep their eyes open and to not fall asleep.

### C. Image preprocessing

All images were preprocessed using the statistical parametric mapping software (SPM12, https://www.fil.ion.ucl.ac.uk/spm/). Briefly, after removing the first 5 volumes, all images were realigned and slice-time corrected. Then, the six rigid motion parameters as well as the white matter and cerebrospinal fluid signals were regressed from all images, which were subsequently normalized to MNI space and band-pass filtered. The mean timeseries of each brain region included in the 200-node Craddock atlas was extracted for each individual.

### D. Lagged correlation

The lagged correlation between the activation time series of two brain regions (*j* and *k*) with activation time courses *x*_*j*_ and *x*_*k*_ respectively, is calculated as the Pearson’s correlation coefficient between *x*_*j*_ and lagged versions of *x*_*k*_ evaluated as a function of the temporal lag. The lag is the number of repetition times by which *x*_*k*_ is shifted with respect to *x*_*j*_ before calculating the correlation. Therefore, the strength of the functional connectivity between the brain regions *j* and *k* at a given delay *d* is calculated as:

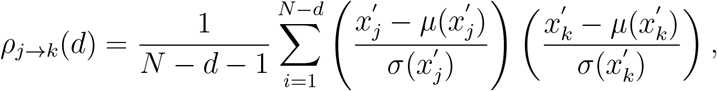

where *N* is the total number of measurements, 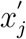 represents the first *N* −*d* measurements of *x*_*j*_ and 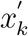 represents the last *N* − *d* measurements of *x*_*k*_; 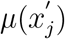 and 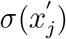 are the mean and standard deviation of 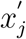 respectively, 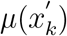 and 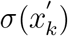 are the mean and standard deviation of 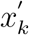.

In this construction, *x*_*k*_ is shifted by *d* time steps with respect to *x*_*j*_, therefore the correlation coefficient *ρ*_*j→k*_(*d*) is an estimation of the directed functional connectivity from region *j* to region *k* due to temporal precedence. By repeating this calculation for all pairs of nodes, we obtain the weighted directed lagged correlation functional network. This network was subsequently binarized at the specified range of densities in order to compare network topologies between the two groups. In this matrix, the directed connection between a pair of nodes *j* and *k* is represented by a pair of elements (*j, k*) and (*k, j*) that quantify the estimated directed influence of brain region *j* to brain region *k* and vice versa.

### E. Anti-symmetric and symmetric correlations

Being a square matrix, the lagged correlation matrix calculated as outlined above can be written as a sum of univocally defined symmetric and anti-symmetric matrices. Therefore, from the lagged correlation matrix *L*, one can calculate the corresponding anti-symmetric matrix *A* as:

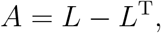

where *L*^T^ denotes the transpose of *L*. As described previously, all negative connections are set to zero. Calculated in this way, the anti-symmetric analysis represents any directed correlation between two regions *j* and *k* with a single entry in the adjacency matrix, which summarizes both the direction and the magnitude of the directed influence.

The symmetric matrix can be calculated as

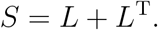

Symmetric matrices do not convey any information about the direction of the functional connections. The magnitude of a connection is calculated as the sum of the connection weights in the directed connections between nodes that run in both directions. The advantage of this method when compared to the zero-lag correlation method is that it can be evaluated at various temporal lags, therefore allowing a more direct comparison with the corresponding directed methods.

### F. Zero-lag correlation method

In the standard zero-lag correlation method, the functional connectivity between two nodes *j* and *k* with respective activation time series *x*_*j*_ and *x*_*k*_ is quantified by the Pearson’s linear correlation coefficient at lag of 0, calculated as:

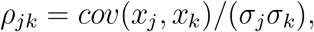

where *cov*(*x*_*j*_, *x*_*k*_) represents the covariance of the corresponding activation time series and *σ*_*j*_ and *σ*_*k*_ are their respective standard deviations. The functional networks are built by calculating the Pearson’s coefficient between all pairs of nodes in the network.

### G. Definition of graph measures

All graph measures were calculated using the Brain Analysis using Graph Theory software [42] (BRAPH, http://braph.org/). In the case of directed binary networks, the indegree of a node is defined as the number of inward edges going into a node. The out-degree of a node is the number of outward edges originating from a node. Denoting the network adjacency matrix with *A* and its elements as *a*_*ij*_, the in- and out-degrees or a node i are expressed as:

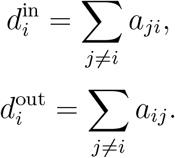

The degree of a node is expressed as the sum of the node’s respective in- and out-degrees:

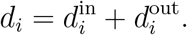

A direct path between two nodes *i* and *j* is the sequence of directed edges that need to be traversed in order to reach *j* starting from *i*. The directed distance 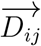 is the number of edges contained in the shortest directed path from *i* to *j*. The regional out-global efficiency of a node *i*, denoted by *e*_out_(*i*), is defined as the average inverse distance from *i* to all other nodes in the network, when considering only directed paths originating from *i*. Analogously, the regional in-global efficiency of node *i, e*_in_(*i*), is the average of the inverse distance to *i* from all other nodes in the network over directed paths ending at *i*. The global counterparts of these measures in a network with *N* nodes can be calculated as the average of the regional out- and in-efficiency of all nodes:

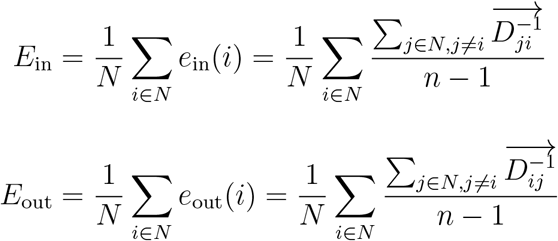

We furthermore calculate the regional in- and out-local efficiency of a node *i* defined as the corresponding global efficiency measure evaluated on the subgraph consisting of nodes that are neighbors of *i*. The in- and out-local efficiency of the network, *LE*_in_ and *LE*_out_ respectively, is calculated by averaging the corresponding measures over all nodes in the network. We define the network’s total global efficiency (*E*) and local efficiency (*LE*) as the mean of the in- and out-efficiency measures:

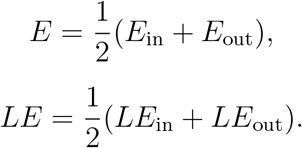

The clustering coefficient *C*_*i*_, of node *i*, reflects the fraction of the neighbors of *i* that are also connected with each other. It can be calculated as the fraction of completed triangles that are present around *i*. In directed networks, we consider a triangle to be completed if its constituent edges form a cycle in either direction. Therefore, we calculate the clustering coefficient as

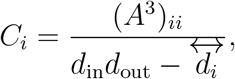

where *d*_in_ and *d*_out_ are the in- and out-degree respectively, and 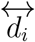 is the number of bilateral edges between i and its neighbors:

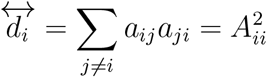

The transitivity indicates the number of triangles present within the complete network. As such, it is calculated as:

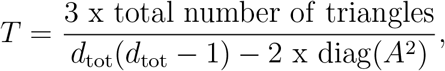

where again we consider a triangle to be completed only if the three directed edges form a cycle, and diag(*A*^2^) is the sum of the diagonal elements in the *A*^2^ matrix.

The modularity quantifies the degree at which a given network can be subdivided into clearly separated communities that have large density of within-community edges and small number of between community edges. Modularity was calculated using Louvain algorithm [10].

### H. Statistical analysis

The statistical significance of the differences between the groups was assessed by performing non-parametric permutation tests with 10000 permutations, which were considered significant for a two-tailed test of the null hypothesis at *p <* 0.05. Additionally, the regional network results were adjusted for multiple comparisons by applying false discovery rate (FDR) corrections at *q <* 0.05 using the Benjamini-Hochberg procedure [5] to control for the number of regions. Non-parametric permutation tests with 10000 permutations were also used to assess between-group differences in demographic variables. All analyses included age, gender and the 6 rigid-body motion parameters as covariates.

## V. ETHICS STATEMENT

The PPMI study was approved by the institutional review board of all participating sites and written informed consent was obtained from all participants before inclusion in the study.

## VII. ACKNOWLEDGEMENTS

Data used in the preparation of this article were obtained from the Parkinson’s Progression Markers Initiative (PPMI) database (www.ppmi-info.org/data). For up-to-date information on the study, visit www.ppmi-info.org. PPMI – a public-private partnership – is funded by the Michael J. Fox Foundation for Parkinson’s Research and funding partners, including Abbvie, Allergan, Amathus, Avid, Biogen, Biolegend, Bristol-Myers-Squibb, Celgene, Denali, GE Healthcare, Genentech, Glaxo-Smith-Kline, Golub Capital, Handl, Insitro, Janssen, Lilly, Lundbeck, Merck, Meso Scale Discovery, Neurocrine, Pfizer, Piramal, Prevail, Roche, Sanofi Genzyme, Servier, Takeda, Teva, UCB, Verily, Voyager.

