## Supplementary material for "Directed brain connectivity identifies widespread functional network changes in Parkinson’s disease": SI Appendix

#### Supplementary information

Mite Mijalkov,[1,*] Giovanni Volpe,[2] and Joana B. Pereira[1,3,*]

[1]*Department of Neurobiology, Care Sciences and Society, Karolinska Institutet, Stockholm, Sweden*

[2]*Department of Physics, Goteborg University, Goteborg, Sweden*

[3]*Memory Research Unit, Department of Clinical Sciences Malmö, Lund University, Lund, Sweden*

### I. DIFFERENCES IN GLOBAL TOPOLOGY

#### A. Lag 2

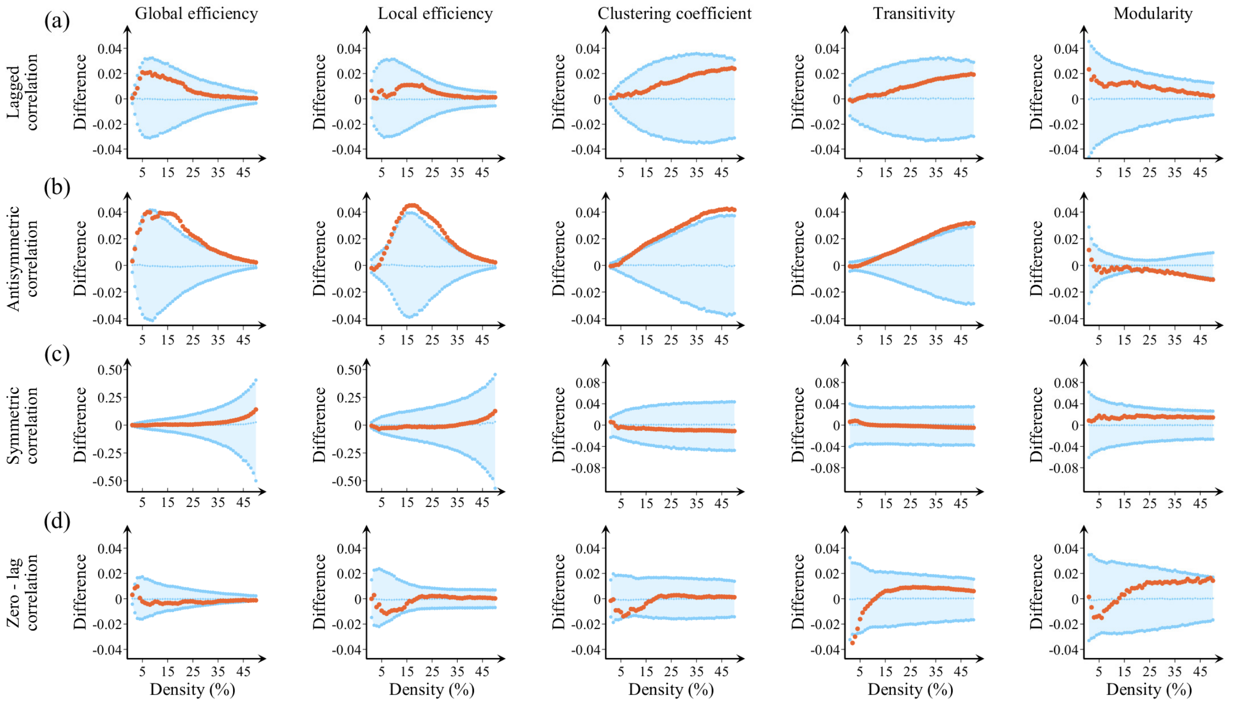


FIG. S1. **Differences between controls and PD patients in global network measures.** Plots showing the differences between controls and PD patients in the global efficiency, local efficiency, clustering coefficient, transitivity and modularity in the case of (a) lagged correlation, (b) anti-symmetric correlation, (c) symmetric correlation and (d) zero-lag correlation. The plots show the upper and lower bounds of the 95% confidence intervals (CI) in blue, and the differences in the network measures between groups in orange circles as a function of network density. The differences are considered statistically significant if they fall outside the CIs.

#### B. Lag 3

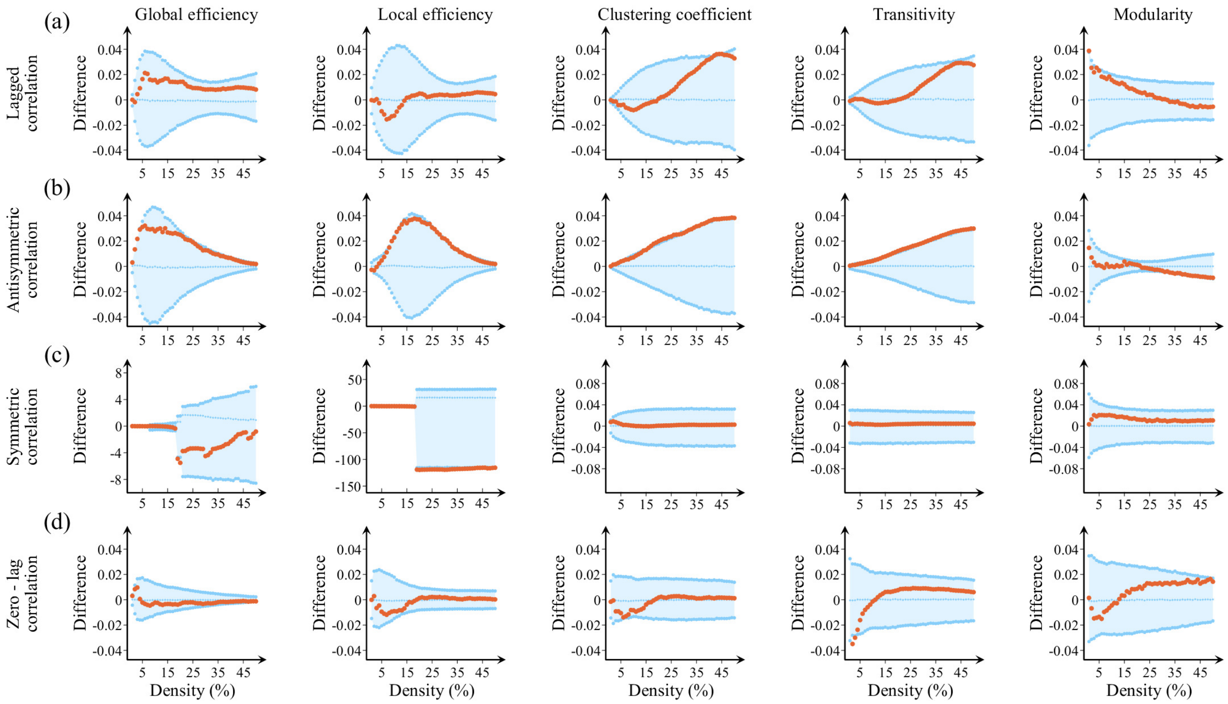


FIG. S2. **Differences between controls and PD patients in global network measures.** Plots showing the differences between controls and PD patients in the global efficiency, local efficiency, clustering coefficient, transitivity and modularity in the case of (a) lagged correlation, (b) anti-symmetric correlation, (c) symmetric correlation and (d) zero-lag correlation. The plots show the upper and lower bounds of the 95% confidence intervals (CI) in blue, and the differences in the network measures between groups in orange circles as a function of network density. The differences are considered statistically significant if they fall outside the CIs.

#### C. Lag 4

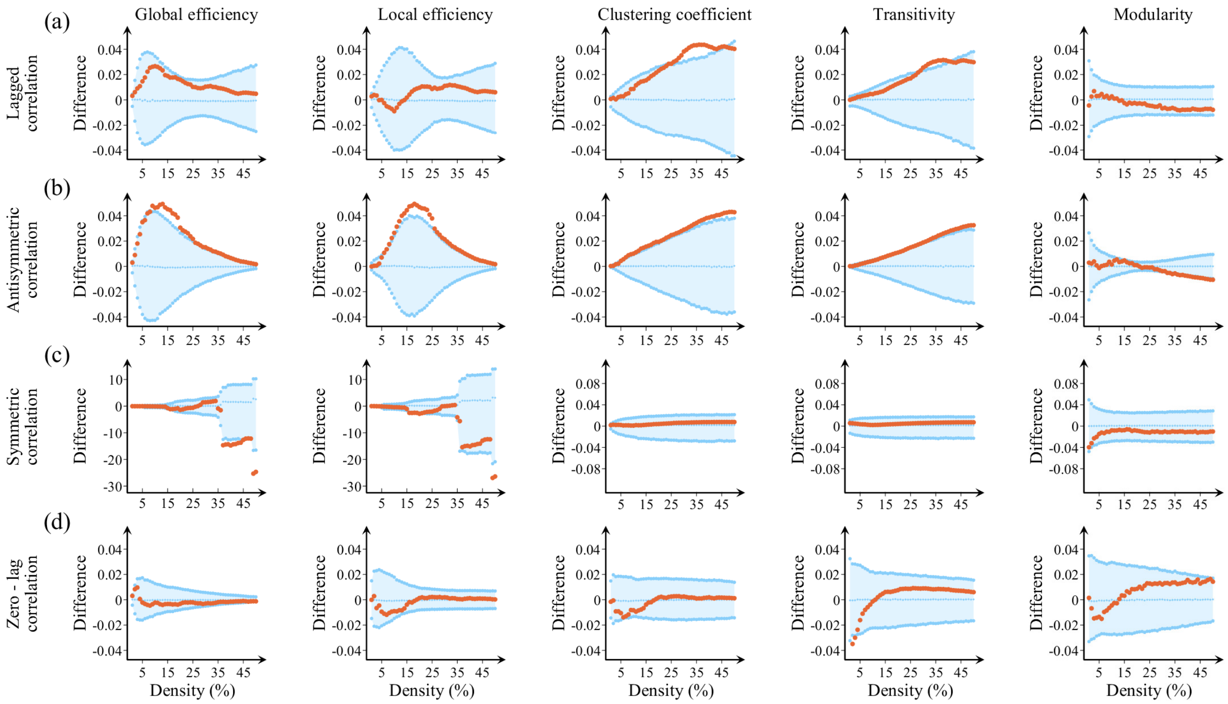


FIG. S3. **Differences between controls and PD patients in global network measures.** Plots showing the differences between controls and PD patients in the global efficiency, local efficiency, clustering coefficient, transitivity and modularity in the case of (a) lagged correlation, (b) anti-symmetric correlation, (c) symmetric correlation and (d) zero-lag correlation. The plots show the upper and lower bounds of the 95% confidence intervals (CI) in blue, and the differences in the network measures between groups in orange circles as a function of network density. The differences are considered statistically significant if they fall outside the CIs.

#### D. Lag 5

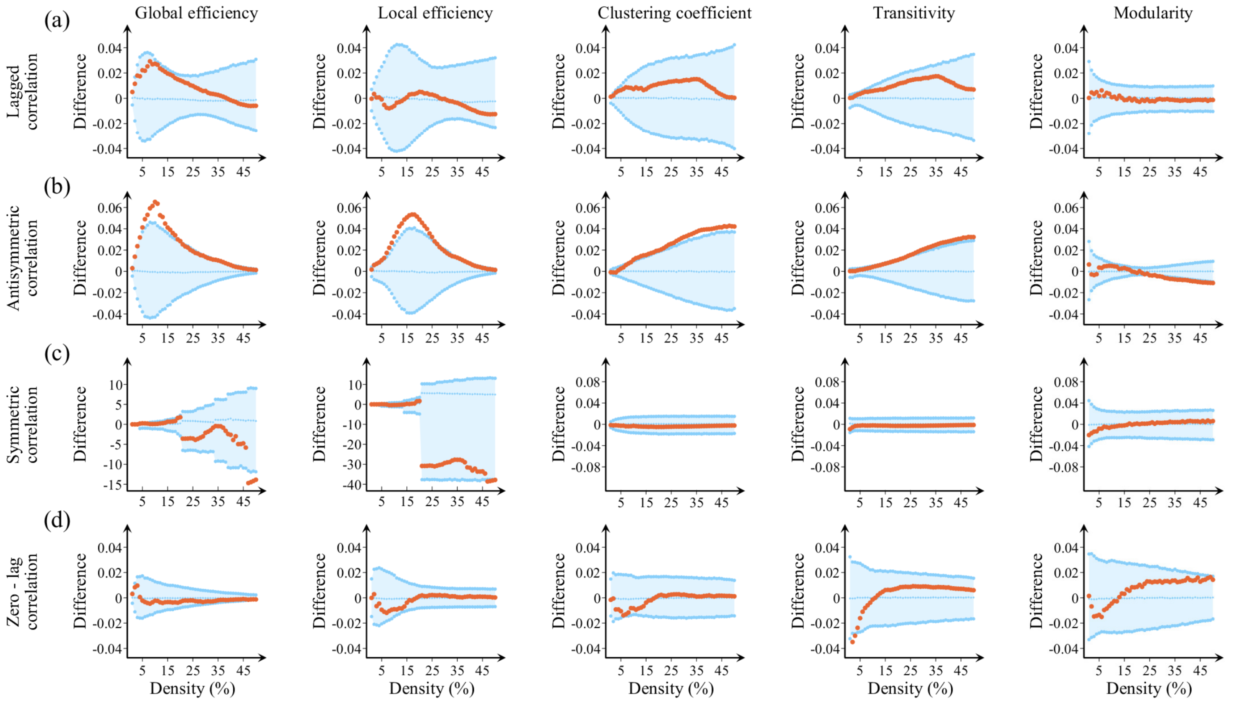


FIG. S4. **Differences between controls and PD patients in global network measures.** Plots showing the differences between controls and PD patients in the global efficiency, local efficiency, clustering coefficient, transitivity and modularity in the case of (a) lagged correlation, (b) anti-symmetric correlation, (c) symmetric correlation and (d) zero-lag correlation. The plots show the upper and lower bounds of the 95% confidence intervals (CI) in blue, and the differences in the network measures between groups in orange circles as a function of network density. The differences are considered statistically significant if they fall outside the CIs.

#### E. Lag 6

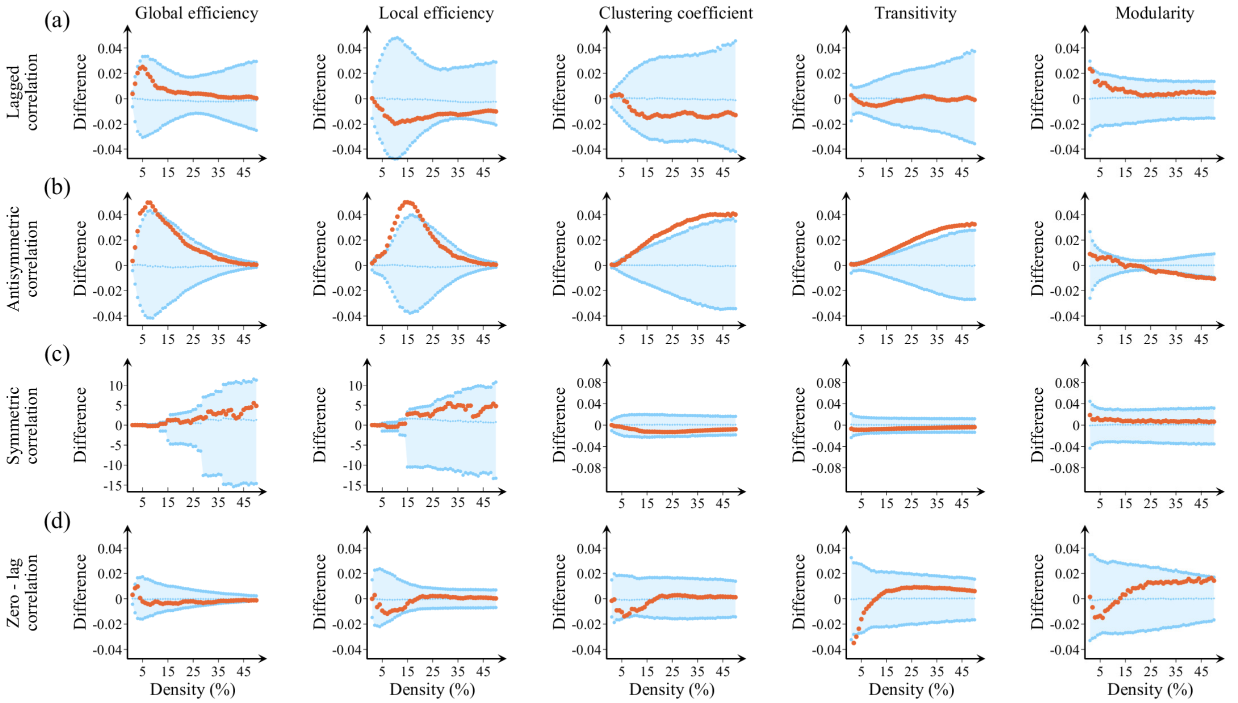


FIG. S5. **Differences between controls and PD patients in global network measures.** Plots showing the differences between controls and PD patients in the global efficiency, local efficiency, clustering coefficient, transitivity and modularity in the case of (a) lagged correlation, (b) anti-symmetric correlation, (c) symmetric correlation and (d) zero-lag correlation. The plots show the upper and lower bounds of the 95% confidence intervals (CI) in blue, and the differences in the network measures between groups in orange circles as a function of network density. The differences are considered statistically significant if they fall outside the CIs.

#### F. Lag 7

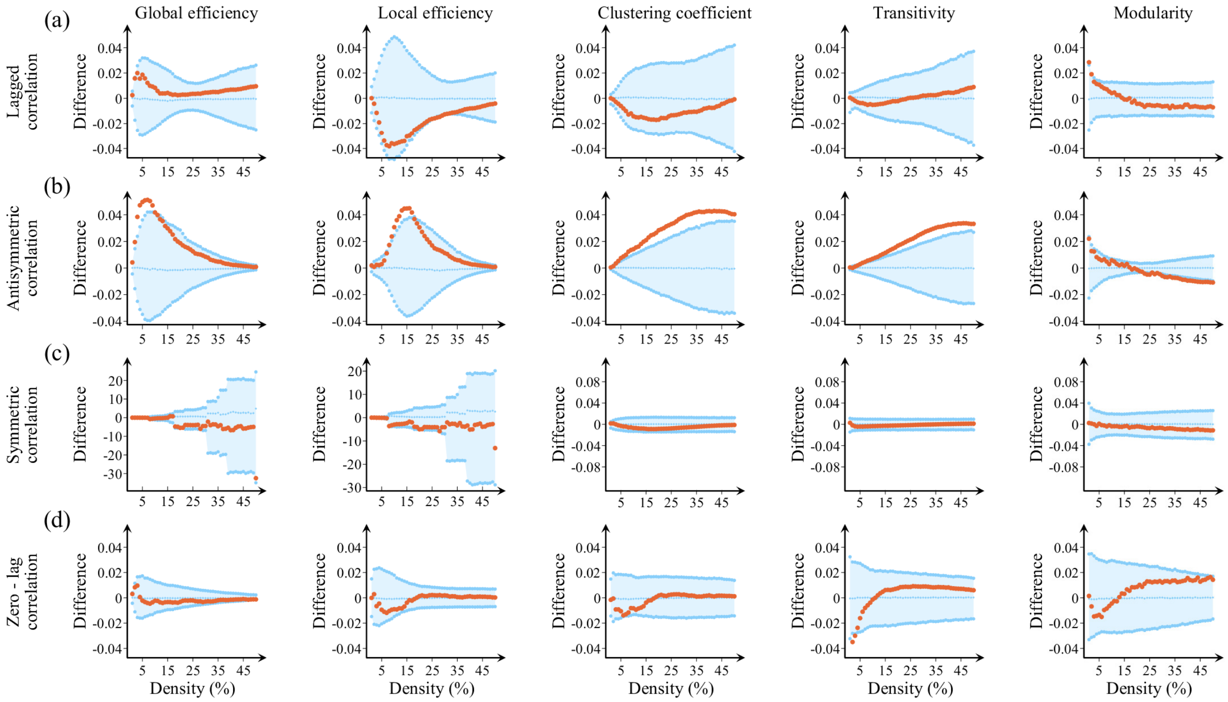


FIG. S6. **Differences between controls and PD patients in global network measures.** Plots showing the differences between controls and PD patients in the global efficiency, local efficiency, clustering coefficient, transitivity and modularity in the case of (a) lagged correlation, (b) anti-symmetric correlation, (c) symmetric correlation and (d) zero-lag correlation. The plots show the upper and lower bounds of the 95% confidence intervals (CI) in blue, and the differences in the network measures between groups in orange circles as a function of network density. The differences are considered statistically significant if they fall outside the CIs.

#### II. CORRELATION ANALYSIS WITH CLINICAL MEASURES IN PD PATIENTS

TABLE S1: Correlation analysis between various global measures calculated at lag of 1 and the UPDRS-III motor scores.

| Global efficiency | | | | | |
|---|---|---|---|---|---|
| **density** | **coefficient** | **p-value** | **density** | **coefficient** | **p-value** |
| 2.000 | 0.330 | 0.001 | 27.000 | 0.279 | 0.007 |
| 3.000 | 0.325 | 0.002 | 28.000 | 0.268 | 0.010 |
| 4.000 | 0.341 | 0.001 | 29.000 | 0.269 | 0.010 |
| 5.000 | 0.343 | 0.001 | 30.000 | 0.270 | 0.010 |
| 6.000 | 0.319 | 0.002 | 31.000 | 0.263 | 0.012 |
| 7.000 | 0.310 | 0.003 | 32.000 | 0.262 | 0.012 |
| 8.000 | 0.297 | 0.004 | 33.000 | 0.257 | 0.014 |
| 9.000 | 0.286 | 0.006 | 34.000 | 0.255 | 0.015 |
| 10.000 | 0.288 | 0.006 | 35.000 | 0.256 | 0.014 |
| 11.000 | 0.283 | 0.007 | 36.000 | 0.259 | 0.013 |
| 12.000 | 0.283 | 0.007 | 37.000 | 0.261 | 0.013 |
| 13.000 | 0.285 | 0.006 | 38.000 | 0.257 | 0.014 |
| 14.000 | 0.298 | 0.004 | 39.000 | 0.254 | 0.015 |
| 15.000 | 0.301 | 0.004 | 40.000 | 0.257 | 0.014 |
| 16.000 | 0.297 | 0.004 | 41.000 | 0.253 | 0.015 |
| 17.000 | 0.291 | 0.005 | 42.000 | 0.252 | 0.016 |
| 18.000 | 0.290 | 0.005 | 43.000 | 0.248 | 0.018 |
| 19.000 | 0.290 | 0.005 | 44.000 | 0.252 | 0.016 |
| 20.000 | 0.286 | 0.006 | 45.000 | 0.238 | 0.023 |
| 21.000 | 0.282 | 0.007 | 46.000 | 0.249 | 0.017 |
| 22.000 | 0.284 | 0.006 | 47.000 | 0.256 | 0.014 |
| 23.000 | 0.275 | 0.008 | 48.000 | 0.255 | 0.015 |
| 24.000 | 0.273 | 0.009 | 49.000 | 0.262 | 0.012 |

Continued on next page

**TABLE S1 – continued from previous page**

| | | | | | |
|---|---|---|---|---|---|
| 25.000 | 0.278 | 0.008 | 50.000 | 0.257 | 0.014 |
| 26.000 | 0.282 | 0.007 | | | |

**Local efficiency**

| density | coefficient | p-value | density | coefficient | p-value |
|---|---|---|---|---|---|
| 10.000 | 0.215 | 0.041 | 31.000 | 0.268 | 0.010 |
| 11.000 | 0.234 | 0.026 | 32.000 | 0.269 | 0.010 |
| 12.000 | 0.245 | 0.019 | 33.000 | 0.264 | 0.011 |
| 13.000 | 0.256 | 0.014 | 34.000 | 0.264 | 0.011 |
| 14.000 | 0.280 | 0.007 | 35.000 | 0.265 | 0.011 |
| 15.000 | 0.290 | 0.005 | 36.000 | 0.268 | 0.010 |
| 16.000 | 0.281 | 0.007 | 37.000 | 0.269 | 0.010 |
| 17.000 | 0.271 | 0.009 | 38.000 | 0.265 | 0.011 |
| 18.000 | 0.272 | 0.009 | 39.000 | 0.258 | 0.013 |
| 19.000 | 0.273 | 0.009 | 40.000 | 0.260 | 0.013 |
| 20.000 | 0.271 | 0.009 | 41.000 | 0.256 | 0.014 |
| 21.000 | 0.267 | 0.011 | 42.000 | 0.255 | 0.015 |
| 22.000 | 0.267 | 0.011 | 43.000 | 0.250 | 0.017 |
| 23.000 | 0.259 | 0.013 | 44.000 | 0.252 | 0.016 |
| 24.000 | 0.258 | 0.014 | 45.000 | 0.239 | 0.022 |
| 25.000 | 0.267 | 0.011 | 46.000 | 0.249 | 0.017 |
| 26.000 | 0.270 | 0.010 | 47.000 | 0.256 | 0.014 |
| 27.000 | 0.268 | 0.010 | 48.000 | 0.254 | 0.015 |
| 28.000 | 0.261 | 0.013 | 49.000 | 0.261 | 0.012 |
| 29.000 | 0.266 | 0.011 | 50.000 | 0.257 | 0.014 |
| 30.000 | 0.270 | 0.010 | | | |

**Clustering coefficient**

| density | coefficient | p-value | density | coefficient | p-value |
|---|---|---|---|---|---|

Continued on next page

**TABLE S1 – continued from previous page**

| | | | | | |
|---|---|---|---|---|---|
| 16.000 | 0.295 | 0.005 | 34.000 | 0.279 | 0.008 |
| 17.000 | 0.288 | 0.006 | 35.000 | 0.277 | 0.008 |
| 18.000 | 0.291 | 0.005 | 36.000 | 0.280 | 0.007 |
| 19.000 | 0.287 | 0.006 | 37.000 | 0.279 | 0.007 |
| 20.000 | 0.284 | 0.006 | 38.000 | 0.278 | 0.008 |
| 21.000 | 0.280 | 0.007 | 39.000 | 0.278 | 0.008 |
| 22.000 | 0.279 | 0.007 | 40.000 | 0.280 | 0.007 |
| 23.000 | 0.277 | 0.008 | 41.000 | 0.279 | 0.007 |
| 24.000 | 0.278 | 0.008 | 42.000 | 0.278 | 0.008 |
| 25.000 | 0.283 | 0.007 | 43.000 | 0.277 | 0.008 |
| 26.000 | 0.283 | 0.007 | 44.000 | 0.278 | 0.008 |
| 27.000 | 0.282 | 0.007 | 45.000 | 0.275 | 0.008 |
| 28.000 | 0.277 | 0.008 | 46.000 | 0.276 | 0.008 |
| 29.000 | 0.280 | 0.007 | 47.000 | 0.277 | 0.008 |
| 30.000 | 0.282 | 0.007 | 48.000 | 0.278 | 0.008 |
| 31.000 | 0.277 | 0.008 | 49.000 | 0.280 | 0.007 |
| 32.000 | 0.278 | 0.008 | 50.000 | 0.279 | 0.007 |
| 33.000 | 0.276 | 0.008 | | | |

**Transitivity**

| **density** | **coefficient** | **p-value** | **density** | **coefficient** | **p-value** |
|---|---|---|---|---|---|
| 20.000 | 0.266 | 0.011 | 36.000 | 0.282 | 0.007 |
| 21.000 | 0.267 | 0.011 | 37.000 | 0.281 | 0.007 |
| 22.000 | 0.269 | 0.010 | 38.000 | 0.280 | 0.007 |
| 23.000 | 0.270 | 0.010 | 39.000 | 0.282 | 0.007 |
| 24.000 | 0.270 | 0.010 | 40.000 | 0.283 | 0.007 |
| 25.000 | 0.276 | 0.008 | 41.000 | 0.282 | 0.007 |
| 26.000 | 0.278 | 0.008 | 42.000 | 0.281 | 0.007 |

Continued on next page

**TABLE S1 – continued from previous page**

| | | | | | |
|---|---|---|---|---|---|
| 27.000 | 0.279 | 0.007 | 43.000 | 0.280 | 0.007 |
| 28.000 | 0.277 | 0.008 | 44.000 | 0.281 | 0.007 |
| 29.000 | 0.280 | 0.007 | 45.000 | 0.278 | 0.008 |
| 30.000 | 0.281 | 0.007 | 46.000 | 0.279 | 0.007 |
| 31.000 | 0.277 | 0.008 | 47.000 | 0.280 | 0.007 |
| 32.000 | 0.278 | 0.008 | 48.000 | 0.281 | 0.007 |
| 33.000 | 0.277 | 0.008 | 49.000 | 0.282 | 0.007 |
| 34.000 | 0.281 | 0.007 | 50.000 | 0.281 | 0.007 |
| 35.000 | 0.281 | 0.007 | | | |

| **Modularity** | | | | | |
|---|---|---|---|---|---|
| **density** | **coefficient** | **p-value** | **density** | **coefficient** | **p-value** |
| 21.000 | -0.248 | 0.018 | 36.000 | -0.231 | 0.028 |
| 22.000 | -0.305 | 0.003 | 37.000 | -0.245 | 0.019 |
| 23.000 | -0.344 | 0.001 | 38.000 | -0.239 | 0.022 |
| 24.000 | -0.265 | 0.011 | 39.000 | -0.261 | 0.013 |
| 25.000 | -0.371 | 0.000 | 40.000 | -0.265 | 0.011 |
| 26.000 | -0.322 | 0.002 | 41.000 | -0.267 | 0.010 |
| 27.000 | -0.323 | 0.002 | 42.000 | -0.271 | 0.009 |
| 28.000 | -0.260 | 0.013 | 43.000 | -0.275 | 0.008 |
| 29.000 | -0.281 | 0.007 | 44.000 | -0.273 | 0.009 |
| 30.000 | -0.319 | 0.002 | 45.000 | -0.272 | 0.009 |
| 31.000 | -0.254 | 0.015 | 46.000 | -0.272 | 0.009 |
| 32.000 | -0.303 | 0.004 | 47.000 | -0.273 | 0.009 |
| 33.000 | -0.263 | 0.012 | 48.000 | -0.277 | 0.008 |
| 34.000 | -0.249 | 0.017 | 49.000 | -0.279 | 0.007 |
| 35.000 | -0.254 | 0.015 | 50.000 | -0.277 | 0.008 |

TABLE S2: Correlation analysis between various global measures calculated at lag of 2 and the UPDRS-III motor scores.

| **Global efficiency** | | | | | |
|---|---|---|---|---|---|
| **density** | **coefficient** | **p-value** | **density** | **coefficient** | **p-value** |
| 12.000 | 0.262 | 0.012 | 32.000 | 0.263 | 0.012 |
| 13.000 | 0.263 | 0.012 | 33.000 | 0.264 | 0.011 |
| 14.000 | 0.264 | 0.011 | 34.000 | 0.257 | 0.014 |
| 15.000 | 0.271 | 0.009 | 35.000 | 0.260 | 0.013 |
| 16.000 | 0.268 | 0.010 | 36.000 | 0.263 | 0.012 |
| 17.000 | 0.266 | 0.011 | 37.000 | 0.258 | 0.014 |
| 18.000 | 0.270 | 0.010 | 38.000 | 0.257 | 0.014 |
| 19.000 | 0.283 | 0.007 | 39.000 | 0.254 | 0.015 |
| 20.000 | 0.271 | 0.009 | 40.000 | 0.251 | 0.016 |
| 21.000 | 0.271 | 0.009 | 41.000 | 0.250 | 0.017 |
| 22.000 | 0.277 | 0.008 | 42.000 | 0.252 | 0.016 |
| 23.000 | 0.280 | 0.007 | 43.000 | 0.248 | 0.018 |
| 24.000 | 0.278 | 0.008 | 44.000 | 0.249 | 0.017 |
| 25.000 | 0.276 | 0.008 | 45.000 | 0.247 | 0.018 |
| 26.000 | 0.275 | 0.008 | 46.000 | 0.243 | 0.020 |
| 27.000 | 0.270 | 0.010 | 47.000 | 0.245 | 0.019 |
| 28.000 | 0.265 | 0.011 | 48.000 | 0.241 | 0.021 |
| 29.000 | 0.264 | 0.011 | 49.000 | 0.237 | 0.024 |
| 30.000 | 0.272 | 0.009 | 50.000 | 0.246 | 0.019 |
| 31.000 | 0.267 | 0.011 | NaN | NaN | NaN |
| **Local efficiency** | | | | | |
| **density** | **coefficient** | **p-value** | **density** | **coefficient** | **p-value** |
| 12.000 | 0.207 | 0.049 | 32.000 | 0.258 | 0.014 |
| 13.000 | 0.218 | 0.038 | 33.000 | 0.263 | 0.012 |

Continued on next page

| | | | | | |
|---|---|---|---|---|---|
| 14.000 | 0.220 | 0.036 | 34.000 | 0.259 | 0.013 |
| 15.000 | 0.229 | 0.029 | 35.000 | 0.260 | 0.013 |
| 16.000 | 0.227 | 0.031 | 36.000 | 0.263 | 0.012 |
| 17.000 | 0.227 | 0.030 | 37.000 | 0.259 | 0.013 |
| 18.000 | 0.227 | 0.031 | 38.000 | 0.257 | 0.014 |
| 19.000 | 0.234 | 0.026 | 39.000 | 0.255 | 0.015 |
| 20.000 | 0.231 | 0.028 | 40.000 | 0.253 | 0.016 |
| 21.000 | 0.231 | 0.028 | 41.000 | 0.253 | 0.016 |
| 22.000 | 0.241 | 0.021 | 42.000 | 0.255 | 0.015 |
| 23.000 | 0.248 | 0.018 | 43.000 | 0.250 | 0.017 |
| 24.000 | 0.253 | 0.016 | 44.000 | 0.251 | 0.017 |
| 25.000 | 0.254 | 0.015 | 45.000 | 0.249 | 0.017 |
| 26.000 | 0.255 | 0.015 | 46.000 | 0.243 | 0.020 |
| 27.000 | 0.254 | 0.015 | 47.000 | 0.244 | 0.020 |
| 28.000 | 0.254 | 0.015 | 48.000 | 0.242 | 0.021 |
| 29.000 | 0.255 | 0.015 | 49.000 | 0.237 | 0.024 |
| 30.000 | 0.263 | 0.012 | 50.000 | 0.246 | 0.019 |
| 31.000 | 0.261 | 0.012 | NaN | NaN | NaN |
| **Clustering coefficient** | | | | | |
| **density** | **coefficient** | **p-value** | **density** | **coefficient** | **p-value** |
| 11.000 | 0.261 | 0.012 | 31.000 | 0.281 | 0.007 |
| 12.000 | 0.269 | 0.010 | 32.000 | 0.279 | 0.007 |
| 13.000 | 0.279 | 0.007 | 33.000 | 0.282 | 0.007 |
| 14.000 | 0.275 | 0.008 | 34.000 | 0.281 | 0.007 |
| 15.000 | 0.265 | 0.011 | 35.000 | 0.282 | 0.007 |
| 16.000 | 0.256 | 0.014 | 36.000 | 0.282 | 0.007 |
| 17.000 | 0.259 | 0.013 | 37.000 | 0.279 | 0.007 |

Continued on next page

**TABLE S2 – continued from previous page**

| | | | | | |
|---|---|---|---|---|---|
| 18.000 | 0.264 | 0.012 | 38.000 | 0.280 | 0.007 |
| 19.000 | 0.261 | 0.012 | 39.000 | 0.281 | 0.007 |
| 20.000 | 0.263 | 0.012 | 40.000 | 0.281 | 0.007 |
| 21.000 | 0.264 | 0.011 | 41.000 | 0.283 | 0.007 |
| 22.000 | 0.265 | 0.011 | 42.000 | 0.285 | 0.006 |
| 23.000 | 0.271 | 0.009 | 43.000 | 0.284 | 0.006 |
| 24.000 | 0.276 | 0.008 | 44.000 | 0.284 | 0.006 |
| 25.000 | 0.277 | 0.008 | 45.000 | 0.284 | 0.006 |
| 26.000 | 0.282 | 0.007 | 46.000 | 0.282 | 0.007 |
| 27.000 | 0.281 | 0.007 | 47.000 | 0.282 | 0.007 |
| 28.000 | 0.279 | 0.007 | 48.000 | 0.283 | 0.007 |
| 29.000 | 0.278 | 0.008 | 49.000 | 0.283 | 0.007 |
| 30.000 | 0.282 | 0.007 | 50.000 | 0.283 | 0.006 |

**Transitivity**

| **density** | **coefficient** | **p-value** | **density** | **coefficient** | **p-value** |
|---|---|---|---|---|---|
| 21.000 | 0.276 | 0.008 | 36.000 | 0.289 | 0.005 |
| 22.000 | 0.277 | 0.008 | 37.000 | 0.288 | 0.006 |
| 23.000 | 0.283 | 0.007 | 38.000 | 0.288 | 0.006 |
| 24.000 | 0.285 | 0.006 | 39.000 | 0.290 | 0.005 |
| 25.000 | 0.286 | 0.006 | 40.000 | 0.290 | 0.005 |
| 26.000 | 0.289 | 0.005 | 41.000 | 0.290 | 0.005 |
| 27.000 | 0.290 | 0.005 | 42.000 | 0.292 | 0.005 |
| 28.000 | 0.286 | 0.006 | 43.000 | 0.291 | 0.005 |
| 29.000 | 0.287 | 0.006 | 44.000 | 0.292 | 0.005 |
| 30.000 | 0.290 | 0.005 | 45.000 | 0.291 | 0.005 |
| 31.000 | 0.289 | 0.006 | 46.000 | 0.289 | 0.005 |
| 32.000 | 0.287 | 0.006 | 47.000 | 0.289 | 0.005 |

Continued on next page

**TABLE S2 – continued from previous page**

| | | | | | |
|---|---|---|---|---|---|
| 33.000 | 0.289 | 0.005 | 48.000 | 0.290 | 0.005 |
| 34.000 | 0.289 | 0.005 | 49.000 | 0.290 | 0.005 |
| 35.000 | 0.290 | 0.005 | 50.000 | 0.290 | 0.005 |

| Modularity | | | | | |
|---|---|---|---|---|---|
| **density** | **coefficient** | **p-value** | **density** | **coefficient** | **p-value** |
| 28.000 | -0.271 | 0.009 | 40.000 | -0.257 | 0.014 |
| 29.000 | -0.277 | 0.008 | 41.000 | -0.266 | 0.011 |
| 30.000 | -0.332 | 0.001 | 42.000 | -0.276 | 0.008 |
| 31.000 | -0.314 | 0.002 | 43.000 | -0.275 | 0.008 |
| 32.000 | -0.295 | 0.005 | 44.000 | -0.275 | 0.008 |
| 33.000 | -0.279 | 0.007 | 45.000 | -0.278 | 0.008 |
| 34.000 | -0.284 | 0.006 | 46.000 | -0.274 | 0.008 |
| 35.000 | -0.267 | 0.010 | 47.000 | -0.275 | 0.008 |
| 36.000 | -0.267 | 0.011 | 48.000 | -0.275 | 0.008 |
| 37.000 | -0.270 | 0.010 | 49.000 | -0.278 | 0.008 |
| 38.000 | -0.255 | 0.015 | 50.000 | -0.280 | 0.007 |
| 39.000 | -0.251 | 0.016 | NaN | NaN | NaN |

TABLE S3: Correlation analysis between various global measures calculated at lag of 3 and the UPDRS-III motor scores.

| **Clustering coefficient** | | | | | |
|---|---|---|---|---|---|
| **density** | **coefficient** | **p-value** | **density** | **coefficient** | **p-value** |
| 16.000 | 0.261 | 0.012 | 38.000 | 0.280 | 0.007 |
| 17.000 | 0.265 | 0.011 | 39.000 | 0.281 | 0.007 |
| 18.000 | 0.264 | 0.012 | 40.000 | 0.283 | 0.007 |
| 19.000 | 0.268 | 0.010 | 41.000 | 0.283 | 0.007 |
| 20.000 | 0.268 | 0.010 | 42.000 | 0.281 | 0.007 |
| 21.000 | 0.267 | 0.010 | 43.000 | 0.281 | 0.007 |
| 22.000 | 0.267 | 0.011 | 44.000 | 0.281 | 0.007 |
| 23.000 | 0.270 | 0.010 | 45.000 | 0.284 | 0.006 |
| 24.000 | 0.272 | 0.009 | 46.000 | 0.286 | 0.006 |
| 25.000 | 0.274 | 0.009 | 47.000 | 0.284 | 0.006 |
| 26.000 | 0.271 | 0.009 | 48.000 | 0.285 | 0.006 |
| 27.000 | 0.270 | 0.010 | 49.000 | 0.284 | 0.006 |
| 28.000 | 0.274 | 0.009 | 50.000 | 0.283 | 0.007 |
| 37.000 | 0.279 | 0.008 | NaN | NaN | NaN |
| **Transitivity** | | | | | |
| **density** | **coefficient** | **p-value** | **density** | **coefficient** | **p-value** |
| 14.000 | 0.248 | 0.018 | 33.000 | 0.309 | 0.003 |
| 15.000 | 0.260 | 0.013 | 34.000 | 0.308 | 0.003 |
| 16.000 | 0.265 | 0.011 | 35.000 | 0.307 | 0.003 |
| 17.000 | 0.272 | 0.009 | 36.000 | 0.304 | 0.003 |
| 18.000 | 0.278 | 0.008 | 37.000 | 0.302 | 0.004 |
| 19.000 | 0.286 | 0.006 | 38.000 | 0.302 | 0.004 |
| 20.000 | 0.288 | 0.006 | 39.000 | 0.303 | 0.004 |
| 21.000 | 0.290 | 0.005 | 40.000 | 0.303 | 0.003 |

Continued on next page

**TABLE S3 – continued from previous page**

| | | | | | |
|---|---|---|---|---|---|
| 22.000 | 0.293 | 0.005 | 41.000 | 0.302 | 0.004 |
| 23.000 | 0.295 | 0.004 | 42.000 | 0.300 | 0.004 |
| 24.000 | 0.299 | 0.004 | 43.000 | 0.299 | 0.004 |
| 25.000 | 0.302 | 0.004 | 44.000 | 0.299 | 0.004 |
| 26.000 | 0.300 | 0.004 | 45.000 | 0.300 | 0.004 |
| 27.000 | 0.300 | 0.004 | 46.000 | 0.301 | 0.004 |
| 28.000 | 0.304 | 0.003 | 47.000 | 0.299 | 0.004 |
| 29.000 | 0.305 | 0.003 | 48.000 | 0.299 | 0.004 |
| 30.000 | 0.307 | 0.003 | 49.000 | 0.298 | 0.004 |
| 31.000 | 0.308 | 0.003 | 50.000 | 0.296 | 0.004 |
| 32.000 | 0.311 | 0.003 | NaN | NaN | NaN |

TABLE S4: Correlation analysis between various global measures calculated at lag of 4 and the UPDRS-III motor scores.

| Global efficiency | | | | | |
|---|---|---|---|---|---|
| **density** | **coefficient** | **p-value** | **density** | **coefficient** | **p-value** |
| 9.000 | 0.292 | 0.005 | 17.000 | 0.243 | 0.020 |
| 10.000 | 0.286 | 0.006 | 18.000 | 0.238 | 0.023 |
| 11.000 | 0.273 | 0.009 | 19.000 | 0.241 | 0.022 |
| 12.000 | 0.286 | 0.006 | 20.000 | 0.241 | 0.021 |
| 13.000 | 0.275 | 0.008 | 21.000 | 0.231 | 0.028 |
| 14.000 | 0.263 | 0.012 | 22.000 | 0.226 | 0.031 |
| 15.000 | 0.250 | 0.017 | 23.000 | 0.218 | 0.038 |
| 16.000 | 0.244 | 0.020 | 25.000 | 0.220 | 0.036 |

| Local efficiency | | | | | |
|---|---|---|---|---|---|
| **density** | **coefficient** | **p-value** | **density** | **coefficient** | **p-value** |
| 10.000 | 0.241 | 0.021 | 16.000 | 0.263 | 0.012 |
| 11.000 | 0.268 | 0.010 | 17.000 | 0.257 | 0.014 |
| 12.000 | 0.279 | 0.007 | 18.000 | 0.251 | 0.016 |
| 13.000 | 0.284 | 0.006 | 19.000 | 0.243 | 0.020 |
| 14.000 | 0.281 | 0.007 | 20.000 | 0.239 | 0.023 |
| 15.000 | 0.269 | 0.010 | 21.000 | 0.223 | 0.034 |

| Clustering | | | | | |
|---|---|---|---|---|---|
| **density** | **coefficient** | **p-value** | **density** | **coefficient** | **p-value** |
| 6.000 | 0.247 | 0.018 | 29.000 | 0.274 | 0.009 |
| 7.000 | 0.270 | 0.010 | 30.000 | 0.273 | 0.009 |
| 8.000 | 0.277 | 0.008 | 31.000 | 0.273 | 0.009 |
| 9.000 | 0.252 | 0.016 | 32.000 | 0.271 | 0.009 |
| 10.000 | 0.261 | 0.012 | 33.000 | 0.275 | 0.008 |
| 11.000 | 0.260 | 0.013 | 34.000 | 0.274 | 0.009 |

Continued on next page

**TABLE S4 – continued from previous page**

| | | | | | |
|---|---|---|---|---|---|
| 12.000 | 0.259 | 0.013 | 35.000 | 0.270 | 0.010 |
| 13.000 | 0.269 | 0.010 | 36.000 | 0.274 | 0.009 |
| 14.000 | 0.264 | 0.012 | 37.000 | 0.272 | 0.009 |
| 15.000 | 0.271 | 0.009 | 38.000 | 0.273 | 0.009 |
| 16.000 | 0.275 | 0.008 | 39.000 | 0.271 | 0.009 |
| 17.000 | 0.272 | 0.009 | 40.000 | 0.270 | 0.010 |
| 18.000 | 0.273 | 0.009 | 41.000 | 0.270 | 0.010 |
| 19.000 | 0.276 | 0.008 | 42.000 | 0.271 | 0.009 |
| 20.000 | 0.277 | 0.008 | 43.000 | 0.266 | 0.011 |
| 21.000 | 0.273 | 0.009 | 44.000 | 0.266 | 0.011 |
| 22.000 | 0.275 | 0.008 | 45.000 | 0.265 | 0.011 |
| 23.000 | 0.275 | 0.008 | 46.000 | 0.263 | 0.012 |
| 24.000 | 0.274 | 0.009 | 47.000 | 0.264 | 0.011 |
| 25.000 | 0.280 | 0.007 | 48.000 | 0.259 | 0.013 |
| 26.000 | 0.279 | 0.007 | 49.000 | 0.264 | 0.012 |
| 27.000 | 0.278 | 0.008 | 50.000 | 0.260 | 0.013 |
| 28.000 | 0.273 | 0.009 | NaN | NaN | NaN |
| **Transitivity** | | | | | |
| **density** | **coefficient** | **p-value** | **density** | **coefficient** | **p-value** |
| 23.000 | 0.302 | 0.004 | 37.000 | 0.293 | 0.005 |
| 24.000 | 0.303 | 0.004 | 38.000 | 0.294 | 0.005 |
| 25.000 | 0.305 | 0.003 | 39.000 | 0.292 | 0.005 |
| 26.000 | 0.306 | 0.003 | 40.000 | 0.291 | 0.005 |
| 27.000 | 0.306 | 0.003 | 41.000 | 0.290 | 0.005 |
| 28.000 | 0.302 | 0.004 | 42.000 | 0.290 | 0.005 |
| 29.000 | 0.302 | 0.004 | 43.000 | 0.286 | 0.006 |
| 30.000 | 0.300 | 0.004 | 44.000 | 0.285 | 0.006 |

Continued on next page

**TABLE S4 – continued from previous page**

| | | | | | |
|---|---|---|---|---|---|
| 31.000 | 0.301 | 0.004 | 45.000 | 0.283 | 0.007 |
| 32.000 | 0.298 | 0.004 | 46.000 | 0.280 | 0.007 |
| 33.000 | 0.300 | 0.004 | 47.000 | 0.280 | 0.007 |
| 34.000 | 0.297 | 0.004 | 48.000 | 0.275 | 0.008 |
| 35.000 | 0.293 | 0.005 | 49.000 | 0.278 | 0.008 |
| 36.000 | 0.295 | 0.005 | 50.000 | 0.275 | 0.008 |

| **Modularity** | | | | | |
|---|---|---|---|---|---|
| **density** | **coefficient** | **p-value** | **density** | **coefficient** | **p-value** |
| 32.000 | -0.335 | 0.001 | 42.000 | -0.287 | 0.006 |
| 33.000 | -0.349 | 0.001 | 43.000 | -0.275 | 0.008 |
| 34.000 | -0.323 | 0.002 | 44.000 | -0.279 | 0.007 |
| 35.000 | -0.309 | 0.003 | 45.000 | -0.277 | 0.008 |
| 36.000 | -0.297 | 0.004 | 46.000 | -0.273 | 0.009 |
| 37.000 | -0.294 | 0.005 | 47.000 | -0.266 | 0.011 |
| 38.000 | -0.276 | 0.008 | 48.000 | -0.261 | 0.012 |
| 39.000 | -0.289 | 0.005 | 49.000 | -0.262 | 0.012 |
| 40.000 | -0.291 | 0.005 | 50.000 | -0.258 | 0.014 |
| 41.000 | -0.290 | 0.005 | NaN | NaN | NaN |

TABLE S5: Correlation analysis between various global measures calculated at lag of 5 and the UPDRS-III motor scores.

| Global efficiency | | | | | |
|---|---|---|---|---|---|
| **density** | **coefficient** | **p-value** | **density** | **coefficient** | **p-value** |
| 3.000 | 0.353 | 0.001 | 12.000 | 0.251 | 0.016 |
| 4.000 | 0.352 | 0.001 | 13.000 | 0.239 | 0.022 |
| 5.000 | 0.343 | 0.001 | 14.000 | 0.229 | 0.029 |
| 6.000 | 0.326 | 0.002 | 15.000 | 0.258 | 0.014 |
| 7.000 | 0.319 | 0.002 | 16.000 | 0.257 | 0.014 |
| 8.000 | 0.302 | 0.004 | 17.000 | 0.261 | 0.013 |
| 9.000 | 0.289 | 0.005 | 18.000 | 0.256 | 0.014 |
| 10.000 | 0.272 | 0.009 | 19.000 | 0.245 | 0.019 |
| 11.000 | 0.261 | 0.012 | 20.000 | 0.246 | 0.019 |
| **Local efficiency** | | | | | |
| **density** | **coefficient** | **p-value** | **density** | **coefficient** | **p-value** |
| 8.000 | 0.224 | 0.033 | 18.000 | 0.279 | 0.007 |
| 9.000 | 0.253 | 0.015 | 19.000 | 0.260 | 0.013 |
| 10.000 | 0.282 | 0.007 | 20.000 | 0.256 | 0.014 |
| 11.000 | 0.303 | 0.003 | 21.000 | 0.250 | 0.017 |
| 12.000 | 0.302 | 0.004 | 22.000 | 0.245 | 0.019 |
| 13.000 | 0.300 | 0.004 | 23.000 | 0.241 | 0.021 |
| 14.000 | 0.302 | 0.004 | 24.000 | 0.234 | 0.026 |
| 15.000 | 0.304 | 0.003 | 25.000 | 0.232 | 0.027 |
| 16.000 | 0.297 | 0.004 | 26.000 | 0.222 | 0.034 |
| 17.000 | 0.293 | 0.005 | 27.000 | 0.220 | 0.036 |
| **Clustering** | | | | | |
| **density** | **coefficient** | **p-value** | **density** | **coefficient** | **p-value** |
| 10.000 | 0.287 | 0.006 | 31.000 | 0.308 | 0.003 |

Continued on next page

**TABLE S5 – continued from previous page**

| | | | | | |
|---|---|---|---|---|---|
| 11.000 | 0.294 | 0.005 | 32.000 | 0.309 | 0.003 |
| 12.000 | 0.277 | 0.008 | 33.000 | 0.310 | 0.003 |
| 13.000 | 0.293 | 0.005 | 34.000 | 0.311 | 0.003 |
| 14.000 | 0.298 | 0.004 | 35.000 | 0.310 | 0.003 |
| 15.000 | 0.303 | 0.004 | 36.000 | 0.310 | 0.003 |
| 16.000 | 0.298 | 0.004 | 37.000 | 0.307 | 0.003 |
| 17.000 | 0.309 | 0.003 | 38.000 | 0.309 | 0.003 |
| 18.000 | 0.304 | 0.003 | 39.000 | 0.308 | 0.003 |
| 19.000 | 0.298 | 0.004 | 40.000 | 0.306 | 0.003 |
| 20.000 | 0.302 | 0.004 | 41.000 | 0.307 | 0.003 |
| 21.000 | 0.305 | 0.003 | 42.000 | 0.305 | 0.003 |
| 22.000 | 0.305 | 0.003 | 43.000 | 0.304 | 0.003 |
| 23.000 | 0.304 | 0.003 | 44.000 | 0.303 | 0.004 |
| 24.000 | 0.306 | 0.003 | 45.000 | 0.302 | 0.004 |
| 25.000 | 0.307 | 0.003 | 46.000 | 0.299 | 0.004 |
| 26.000 | 0.305 | 0.003 | 47.000 | 0.301 | 0.004 |
| 27.000 | 0.309 | 0.003 | 48.000 | 0.304 | 0.003 |
| 28.000 | 0.310 | 0.003 | 49.000 | 0.301 | 0.004 |
| 29.000 | 0.310 | 0.003 | 50.000 | 0.298 | 0.004 |
| 30.000 | 0.308 | 0.003 | NaN | NaN | NaN |

**Transitivity**

| density | coefficient | p-value | density | coefficient | p-value |
|---|---|---|---|---|---|
| 23.000 | 0.353 | 0.001 | 37.000 | 0.335 | 0.001 |
| 24.000 | 0.352 | 0.001 | 38.000 | 0.337 | 0.001 |
| 25.000 | 0.353 | 0.001 | 39.000 | 0.334 | 0.001 |
| 26.000 | 0.351 | 0.001 | 40.000 | 0.332 | 0.001 |
| 27.000 | 0.352 | 0.001 | 41.000 | 0.330 | 0.001 |

Continued on next page

**TABLE S5 – continued from previous page**

| | | | | | |
|---|---|---|---|---|---|
| 28.000 | 0.351 | 0.001 | 42.000 | 0.327 | 0.002 |
| 29.000 | 0.350 | 0.001 | 43.000 | 0.325 | 0.002 |
| 30.000 | 0.348 | 0.001 | 44.000 | 0.323 | 0.002 |
| 31.000 | 0.346 | 0.001 | 45.000 | 0.322 | 0.002 |
| 32.000 | 0.346 | 0.001 | 46.000 | 0.318 | 0.002 |
| 33.000 | 0.344 | 0.001 | 47.000 | 0.320 | 0.002 |
| 34.000 | 0.344 | 0.001 | 48.000 | 0.321 | 0.002 |
| 35.000 | 0.341 | 0.001 | 49.000 | 0.318 | 0.002 |
| 36.000 | 0.340 | 0.001 | 50.000 | 0.315 | 0.002 |
| **Modularity** | | | | | |
| **density** | **coefficient** | **p-value** | **density** | **coefficient** | **p-value** |
| 25.000 | -0.304 | 0.003 | 38.000 | -0.340 | 0.001 |
| 26.000 | -0.286 | 0.006 | 39.000 | -0.326 | 0.002 |
| 27.000 | -0.270 | 0.010 | 40.000 | -0.315 | 0.002 |
| 28.000 | -0.273 | 0.009 | 41.000 | -0.305 | 0.003 |
| 29.000 | -0.308 | 0.003 | 42.000 | -0.306 | 0.003 |
| 30.000 | -0.307 | 0.003 | 43.000 | -0.313 | 0.003 |
| 31.000 | -0.363 | 0.000 | 44.000 | -0.312 | 0.003 |
| 32.000 | -0.360 | 0.000 | 45.000 | -0.308 | 0.003 |
| 33.000 | -0.305 | 0.003 | 46.000 | -0.311 | 0.003 |
| 34.000 | -0.340 | 0.001 | 47.000 | -0.312 | 0.003 |
| 35.000 | -0.340 | 0.001 | 48.000 | -0.313 | 0.002 |
| 36.000 | -0.347 | 0.001 | 49.000 | -0.313 | 0.003 |
| 37.000 | -0.316 | 0.002 | 50.000 | -0.307 | 0.003 |

TABLE S6: Correlation analysis between various global measures calculated at lag of 6 and the UPDRS-III motor scores.

| Global efficiency | | | | | |
|---|---|---|---|---|---|
| **density** | **coefficient** | **p-value** | **density** | **coefficient** | **p-value** |
| 1.000 | 0.307 | 0.003 | 6.000 | 0.357 | 0.001 |
| 2.000 | 0.368 | 0.000 | 7.000 | 0.353 | 0.001 |
| 3.000 | 0.392 | 0.000 | 8.000 | 0.347 | 0.001 |
| 4.000 | 0.368 | 0.000 | 9.000 | 0.331 | 0.001 |
| 5.000 | 0.363 | 0.000 | 10.000 | 0.321 | 0.002 |
| **Local efficiency** | | | | | |
| **density** | **coefficient** | **p-value** | **density** | **coefficient** | **p-value** |
| 7.000 | 0.255 | 0.015 | 14.000 | 0.342 | 0.001 |
| 8.000 | 0.308 | 0.003 | 15.000 | 0.338 | 0.001 |
| 9.000 | 0.335 | 0.001 | 16.000 | 0.327 | 0.002 |
| 10.000 | 0.342 | 0.001 | 17.000 | 0.317 | 0.002 |
| 11.000 | 0.344 | 0.001 | 18.000 | 0.306 | 0.003 |
| 12.000 | 0.341 | 0.001 | 19.000 | 0.299 | 0.004 |
| 13.000 | 0.353 | 0.001 | 20.000 | 0.291 | 0.005 |
| **Clustering** | | | | | |
| **density** | **coefficient** | **p-value** | **density** | **coefficient** | **p-value** |
| 9.000 | 0.325 | 0.002 | 30.000 | 0.332 | 0.001 |
| 10.000 | 0.316 | 0.002 | 31.000 | 0.329 | 0.001 |
| 11.000 | 0.315 | 0.002 | 32.000 | 0.330 | 0.001 |
| 12.000 | 0.313 | 0.003 | 33.000 | 0.329 | 0.001 |
| 13.000 | 0.318 | 0.002 | 34.000 | 0.330 | 0.001 |
| 14.000 | 0.320 | 0.002 | 35.000 | 0.326 | 0.002 |
| 15.000 | 0.325 | 0.002 | 36.000 | 0.328 | 0.001 |
| 16.000 | 0.326 | 0.002 | 37.000 | 0.328 | 0.001 |

Continued on next page

**TABLE S6 – continued from previous page**

| | | | | | |
|---|---|---|---|---|---|
| 17.000 | 0.329 | 0.001 | 38.000 | 0.329 | 0.001 |
| 18.000 | 0.325 | 0.002 | 39.000 | 0.328 | 0.001 |
| 19.000 | 0.329 | 0.001 | 40.000 | 0.326 | 0.002 |
| 20.000 | 0.329 | 0.001 | 41.000 | 0.324 | 0.002 |
| 21.000 | 0.334 | 0.001 | 42.000 | 0.325 | 0.002 |
| 22.000 | 0.331 | 0.001 | 43.000 | 0.326 | 0.002 |
| 23.000 | 0.332 | 0.001 | 44.000 | 0.322 | 0.002 |
| 24.000 | 0.330 | 0.001 | 45.000 | 0.322 | 0.002 |
| 25.000 | 0.328 | 0.002 | 46.000 | 0.324 | 0.002 |
| 26.000 | 0.332 | 0.001 | 47.000 | 0.322 | 0.002 |
| 27.000 | 0.330 | 0.001 | 48.000 | 0.323 | 0.002 |
| 28.000 | 0.335 | 0.001 | 49.000 | 0.326 | 0.002 |
| 29.000 | 0.335 | 0.001 | 50.000 | 0.324 | 0.002 |
| **Transitivity** | | | | | |
| **density** | **coefficient** | **p-value** | **density** | **coefficient** | **p-value** |
| 9.000 | 0.348 | 0.001 | 30.000 | 0.341 | 0.001 |
| 10.000 | 0.350 | 0.001 | 31.000 | 0.339 | 0.001 |
| 11.000 | 0.348 | 0.001 | 32.000 | 0.341 | 0.001 |
| 12.000 | 0.345 | 0.001 | 33.000 | 0.339 | 0.001 |
| 13.000 | 0.343 | 0.001 | 34.000 | 0.339 | 0.001 |
| 14.000 | 0.342 | 0.001 | 35.000 | 0.335 | 0.001 |
| 15.000 | 0.344 | 0.001 | 36.000 | 0.337 | 0.001 |
| 16.000 | 0.347 | 0.001 | 37.000 | 0.337 | 0.001 |
| 17.000 | 0.346 | 0.001 | 38.000 | 0.338 | 0.001 |
| 18.000 | 0.347 | 0.001 | 39.000 | 0.337 | 0.001 |
| 19.000 | 0.347 | 0.001 | 40.000 | 0.336 | 0.001 |
| 20.000 | 0.346 | 0.001 | 41.000 | 0.335 | 0.001 |

Continued on next page

**TABLE S6 – continued from previous page**

| | | | | | |
|---|---|---|---|---|---|
| 21.000 | 0.349 | 0.001 | 42.000 | 0.335 | 0.001 |
| 22.000 | 0.346 | 0.001 | 43.000 | 0.336 | 0.001 |
| 23.000 | 0.347 | 0.001 | 44.000 | 0.332 | 0.001 |
| 24.000 | 0.345 | 0.001 | 45.000 | 0.332 | 0.001 |
| 25.000 | 0.342 | 0.001 | 46.000 | 0.334 | 0.001 |
| 26.000 | 0.344 | 0.001 | 47.000 | 0.333 | 0.001 |
| 27.000 | 0.341 | 0.001 | 48.000 | 0.334 | 0.001 |
| 28.000 | 0.344 | 0.001 | 49.000 | 0.337 | 0.001 |
| 29.000 | 0.343 | 0.001 | 50.000 | 0.335 | 0.001 |

| **Modularity** | | | | | |
|---|---|---|---|---|---|
| **density** | **coefficient** | **p-value** | **density** | **coefficient** | **p-value** |
| 25.000 | -0.312 | 0.003 | 38.000 | -0.325 | 0.002 |
| 26.000 | -0.337 | 0.001 | 39.000 | -0.326 | 0.002 |
| 27.000 | -0.376 | 0.000 | 40.000 | -0.310 | 0.003 |
| 28.000 | -0.295 | 0.005 | 41.000 | -0.308 | 0.003 |
| 29.000 | -0.278 | 0.008 | 42.000 | -0.316 | 0.002 |
| 30.000 | -0.347 | 0.001 | 43.000 | -0.316 | 0.002 |
| 31.000 | -0.286 | 0.006 | 44.000 | -0.308 | 0.003 |
| 32.000 | -0.286 | 0.006 | 45.000 | -0.314 | 0.002 |
| 33.000 | -0.312 | 0.003 | 46.000 | -0.319 | 0.002 |
| 34.000 | -0.294 | 0.005 | 47.000 | -0.321 | 0.002 |
| 35.000 | -0.284 | 0.006 | 48.000 | -0.329 | 0.001 |
| 36.000 | -0.312 | 0.003 | 49.000 | -0.329 | 0.001 |
| 37.000 | -0.311 | 0.003 | 50.000 | -0.328 | 0.001 |

TABLE S7: Correlation analysis between various global measures calculated at lag of 7 and the UPDRS-III motor scores.

| **Global efficiency** | | | | | |
|---|---|---|---|---|---|
| **density** | **coefficient** | **p-value** | **density** | **coefficient** | **p-value** |
| 2.000 | 0.280 | 0.007 | 7.000 | 0.263 | 0.012 |
| 3.000 | 0.301 | 0.004 | 8.000 | 0.263 | 0.012 |
| 4.000 | 0.313 | 0.003 | 9.000 | 0.239 | 0.022 |
| 5.000 | 0.310 | 0.003 | 10.000 | 0.246 | 0.019 |
| 6.000 | 0.287 | 0.006 | 11.000 | 0.233 | 0.026 |
| **Local efficiency** | | | | | |
| **density** | **coefficient** | **p-value** | **density** | **coefficient** | **p-value** |
| 8.000 | 0.213 | 0.042 | 13.000 | 0.262 | 0.012 |
| 9.000 | 0.232 | 0.027 | 14.000 | 0.264 | 0.012 |
| 10.000 | 0.259 | 0.013 | 15.000 | 0.252 | 0.016 |
| 11.000 | 0.268 | 0.010 | 16.000 | 0.246 | 0.019 |
| 12.000 | 0.266 | 0.011 | 17.000 | 0.240 | 0.022 |
| **Clustering** | | | | | |
| **density** | **coefficient** | **p-value** | **density** | **coefficient** | **p-value** |
| 6.000 | 0.238 | 0.023 | 29.000 | 0.286 | 0.006 |
| 7.000 | 0.241 | 0.022 | 30.000 | 0.287 | 0.006 |
| 8.000 | 0.267 | 0.010 | 31.000 | 0.288 | 0.006 |
| 9.000 | 0.261 | 0.013 | 32.000 | 0.287 | 0.006 |
| 10.000 | 0.266 | 0.011 | 33.000 | 0.288 | 0.006 |
| 11.000 | 0.269 | 0.010 | 34.000 | 0.287 | 0.006 |
| 12.000 | 0.266 | 0.011 | 35.000 | 0.288 | 0.006 |
| 13.000 | 0.274 | 0.009 | 36.000 | 0.286 | 0.006 |
| 14.000 | 0.269 | 0.010 | 37.000 | 0.283 | 0.007 |
| 15.000 | 0.261 | 0.012 | 38.000 | 0.285 | 0.006 |

Continued on next page

**TABLE S7 – continued from previous page**

| | | | | | |
|---|---|---|---|---|---|
| 16.000 | 0.269 | 0.010 | 39.000 | 0.285 | 0.006 |
| 17.000 | 0.261 | 0.013 | 40.000 | 0.286 | 0.006 |
| 18.000 | 0.269 | 0.010 | 41.000 | 0.285 | 0.006 |
| 19.000 | 0.274 | 0.009 | 42.000 | 0.283 | 0.007 |
| 20.000 | 0.282 | 0.007 | 43.000 | 0.286 | 0.006 |
| 21.000 | 0.287 | 0.006 | 44.000 | 0.283 | 0.007 |
| 22.000 | 0.287 | 0.006 | 45.000 | 0.287 | 0.006 |
| 23.000 | 0.287 | 0.006 | 46.000 | 0.283 | 0.007 |
| 24.000 | 0.286 | 0.006 | 47.000 | 0.283 | 0.007 |
| 25.000 | 0.287 | 0.006 | 48.000 | 0.283 | 0.007 |
| 26.000 | 0.288 | 0.006 | 49.000 | 0.283 | 0.006 |
| 27.000 | 0.283 | 0.007 | 50.000 | 0.283 | 0.007 |
| 28.000 | 0.287 | 0.006 | NaN | NaN | NaN |
| **Transitivity** | | | | | |
| **density** | **coefficient** | **p-value** | **density** | **coefficient** | **p-value** |
| 10.000 | 0.212 | 0.043 | 31.000 | 0.287 | 0.006 |
| 11.000 | 0.218 | 0.038 | 32.000 | 0.287 | 0.006 |
| 12.000 | 0.216 | 0.039 | 33.000 | 0.289 | 0.005 |
| 13.000 | 0.231 | 0.028 | 34.000 | 0.289 | 0.005 |
| 14.000 | 0.239 | 0.022 | 35.000 | 0.291 | 0.005 |
| 15.000 | 0.237 | 0.024 | 36.000 | 0.290 | 0.005 |
| 16.000 | 0.248 | 0.018 | 37.000 | 0.287 | 0.006 |
| 17.000 | 0.247 | 0.018 | 38.000 | 0.290 | 0.005 |
| 18.000 | 0.255 | 0.015 | 39.000 | 0.290 | 0.005 |
| 19.000 | 0.263 | 0.012 | 40.000 | 0.292 | 0.005 |
| 20.000 | 0.272 | 0.009 | 41.000 | 0.291 | 0.005 |
| 21.000 | 0.277 | 0.008 | 42.000 | 0.289 | 0.005 |

Continued on next page

**TABLE S7 – continued from previous page**

| | | | | | |
|---|---|---|---|---|---|
| 22.000 | 0.278 | 0.008 | 43.000 | 0.293 | 0.005 |
| 23.000 | 0.278 | 0.008 | 44.000 | 0.290 | 0.005 |
| 24.000 | 0.278 | 0.008 | 45.000 | 0.294 | 0.005 |
| 25.000 | 0.281 | 0.007 | 46.000 | 0.291 | 0.005 |
| 26.000 | 0.282 | 0.007 | 47.000 | 0.291 | 0.005 |
| 27.000 | 0.279 | 0.007 | 48.000 | 0.292 | 0.005 |
| 28.000 | 0.285 | 0.006 | 49.000 | 0.293 | 0.005 |
| 29.000 | 0.285 | 0.006 | 50.000 | 0.293 | 0.005 |
| 30.000 | 0.286 | 0.006 | NaN | NaN | NaN |

| **Modularity** | | | | | |
|---|---|---|---|---|---|
| **density** | **coefficient** | **p-value** | **density** | **coefficient** | **p-value** |
| 23.000 | -0.312 | 0.003 | 37.000 | -0.301 | 0.004 |
| 24.000 | -0.411 | 0.000 | 38.000 | -0.303 | 0.004 |
| 25.000 | -0.383 | 0.000 | 39.000 | -0.316 | 0.002 |
| 26.000 | -0.367 | 0.000 | 40.000 | -0.310 | 0.003 |
| 27.000 | -0.340 | 0.001 | 41.000 | -0.300 | 0.004 |
| 28.000 | -0.352 | 0.001 | 42.000 | -0.293 | 0.005 |
| 29.000 | -0.372 | 0.000 | 43.000 | -0.295 | 0.005 |
| 30.000 | -0.375 | 0.000 | 44.000 | -0.296 | 0.004 |
| 31.000 | -0.351 | 0.001 | 45.000 | -0.295 | 0.005 |
| 32.000 | -0.334 | 0.001 | 46.000 | -0.292 | 0.005 |
| 33.000 | -0.331 | 0.001 | 47.000 | -0.291 | 0.005 |
| 34.000 | -0.322 | 0.002 | 48.000 | -0.291 | 0.005 |
| 35.000 | -0.318 | 0.002 | 49.000 | -0.295 | 0.005 |
| 36.000 | -0.307 | 0.003 | 50.000 | -0.295 | 0.005 |

TABLE S8: Correlation analysis between various global measures calculated at lag of 1 and the letter number sequencing test scores.

| Transitivity | | | | | |
|---|---|---|---|---|---|
| **density** | **coefficient** | **p-value** | **density** | **coefficient** | **p-value** |
| 20.000 | -0.229 | 0.029 | 36.000 | -0.248 | 0.018 |
| 21.000 | -0.230 | 0.028 | 37.000 | -0.247 | 0.018 |
| 22.000 | -0.234 | 0.026 | 38.000 | -0.248 | 0.018 |
| 23.000 | -0.236 | 0.024 | 39.000 | -0.249 | 0.017 |
| 24.000 | -0.234 | 0.025 | 40.000 | -0.250 | 0.017 |
| 25.000 | -0.239 | 0.023 | 41.000 | -0.247 | 0.018 |
| 26.000 | -0.237 | 0.024 | 42.000 | -0.246 | 0.019 |
| 27.000 | -0.239 | 0.023 | 43.000 | -0.245 | 0.019 |
| 28.000 | -0.241 | 0.022 | 44.000 | -0.244 | 0.020 |
| 29.000 | -0.242 | 0.021 | 45.000 | -0.242 | 0.021 |
| 30.000 | -0.242 | 0.021 | 46.000 | -0.242 | 0.021 |
| 31.000 | -0.245 | 0.019 | 47.000 | -0.242 | 0.021 |
| 32.000 | -0.244 | 0.020 | 48.000 | -0.243 | 0.020 |
| 33.000 | -0.248 | 0.018 | 49.000 | -0.244 | 0.020 |
| 34.000 | -0.246 | 0.019 | 50.000 | -0.245 | 0.019 |
| 35.000 | -0.248 | 0.018 | NaN | NaN | NaN |

TABLE S9: Correlation analysis between various global measures calculated at lag of 2 and the letter number sequencing test scores.

| **Global efficiency** | | | | | |
|---|---|---|---|---|---|
| **density** | **coefficient** | **p-value** | **density** | **coefficient** | **p-value** |
| 12.000 | -0.259 | 0.013 | 25.000 | -0.249 | 0.017 |
| 13.000 | -0.264 | 0.011 | 26.000 | -0.245 | 0.019 |
| 14.000 | -0.260 | 0.013 | 27.000 | -0.242 | 0.021 |
| 15.000 | -0.262 | 0.012 | 28.000 | -0.247 | 0.018 |
| 16.000 | -0.263 | 0.012 | 29.000 | -0.247 | 0.018 |
| 17.000 | -0.266 | 0.011 | 30.000 | -0.240 | 0.022 |
| 18.000 | -0.262 | 0.012 | 31.000 | -0.238 | 0.023 |
| 19.000 | -0.263 | 0.012 | 32.000 | -0.234 | 0.026 |
| 20.000 | -0.265 | 0.011 | 33.000 | -0.233 | 0.026 |
| 21.000 | -0.265 | 0.011 | 34.000 | -0.234 | 0.025 |
| 22.000 | -0.265 | 0.011 | 35.000 | -0.228 | 0.030 |
| 23.000 | -0.257 | 0.014 | 36.000 | -0.228 | 0.030 |
| 24.000 | -0.257 | 0.014 | 37.000 | -0.228 | 0.030 |
| **Local efficiency** | | | | | |
| **density** | **coefficient** | **p-value** | **density** | **coefficient** | **p-value** |
| 11.000 | -0.235 | 0.025 | 25.000 | -0.258 | 0.013 |
| 12.000 | -0.247 | 0.018 | 26.000 | -0.252 | 0.016 |
| 13.000 | -0.274 | 0.009 | 27.000 | -0.248 | 0.018 |
| 14.000 | -0.276 | 0.008 | 28.000 | -0.249 | 0.017 |
| 15.000 | -0.287 | 0.006 | 29.000 | -0.250 | 0.017 |
| 16.000 | -0.284 | 0.006 | 30.000 | -0.239 | 0.023 |
| 17.000 | -0.287 | 0.006 | 31.000 | -0.235 | 0.025 |
| 18.000 | -0.286 | 0.006 | 32.000 | -0.232 | 0.027 |
| 19.000 | -0.285 | 0.006 | 33.000 | -0.231 | 0.027 |

Continued on next page

**TABLE S9 – continued from previous page**

| | | | | | |
|---|---|---|---|---|---|
| 20.000 | -0.286 | 0.006 | 34.000 | -0.231 | 0.027 |
| 21.000 | -0.285 | 0.006 | 35.000 | -0.228 | 0.030 |
| 22.000 | -0.280 | 0.007 | 36.000 | -0.227 | 0.030 |
| 23.000 | -0.270 | 0.010 | 37.000 | -0.227 | 0.031 |
| 24.000 | -0.265 | 0.011 | NaN | NaN | NaN |
| **Clustering** | | | | | |
| **density** | **coefficient** | **p-value** | **density** | **coefficient** | **p-value** |
| 28.000 | -0.231 | 0.028 | 40.000 | -0.251 | 0.016 |
| 29.000 | -0.231 | 0.028 | 41.000 | -0.252 | 0.016 |
| 30.000 | -0.230 | 0.028 | 42.000 | -0.256 | 0.014 |
| 31.000 | -0.234 | 0.026 | 43.000 | -0.257 | 0.014 |
| 32.000 | -0.235 | 0.025 | 44.000 | -0.257 | 0.014 |
| 33.000 | -0.237 | 0.024 | 45.000 | -0.259 | 0.013 |
| 34.000 | -0.239 | 0.022 | 46.000 | -0.260 | 0.013 |
| 35.000 | -0.240 | 0.022 | 47.000 | -0.263 | 0.012 |
| 36.000 | -0.244 | 0.020 | 48.000 | -0.265 | 0.011 |
| 37.000 | -0.246 | 0.019 | 49.000 | -0.266 | 0.011 |
| 38.000 | -0.248 | 0.018 | 50.000 | -0.265 | 0.011 |
| 39.000 | -0.249 | 0.017 | NaN | NaN | NaN |
| **Transitivity** | | | | | |
| **density** | **coefficient** | **p-value** | **density** | **coefficient** | **p-value** |
| 22.000 | -0.208 | 0.048 | 37.000 | -0.249 | 0.017 |
| 23.000 | -0.211 | 0.045 | 38.000 | -0.251 | 0.017 |
| 24.000 | -0.213 | 0.042 | 39.000 | -0.253 | 0.015 |
| 25.000 | -0.219 | 0.037 | 40.000 | -0.256 | 0.014 |
| 26.000 | -0.223 | 0.034 | 41.000 | -0.257 | 0.014 |
| 27.000 | -0.226 | 0.031 | 42.000 | -0.261 | 0.012 |

Continued on next page

**TABLE S9 – continued from previous page**

| | | | | | |
|---|---|---|---|---|---|
| 28.000 | -0.230 | 0.028 | 43.000 | -0.261 | 0.012 |
| 29.000 | -0.232 | 0.027 | 44.000 | -0.261 | 0.012 |
| 30.000 | -0.234 | 0.026 | 45.000 | -0.263 | 0.012 |
| 31.000 | -0.237 | 0.024 | 46.000 | -0.263 | 0.012 |
| 32.000 | -0.238 | 0.023 | 47.000 | -0.266 | 0.011 |
| 33.000 | -0.240 | 0.022 | 48.000 | -0.268 | 0.010 |
| 34.000 | -0.243 | 0.020 | 49.000 | -0.269 | 0.010 |
| 35.000 | -0.245 | 0.019 | 50.000 | -0.268 | 0.010 |
| 36.000 | -0.247 | 0.018 | NaN | NaN | NaN |
| **Modularity** | | | | | |
| **density** | **coefficient** | **p-value** | **density** | **coefficient** | **p-value** |
| 34.000 | 0.236 | 0.024 | 43.000 | 0.266 | 0.011 |
| 35.000 | 0.219 | 0.037 | 44.000 | 0.268 | 0.010 |
| 36.000 | 0.238 | 0.023 | 45.000 | 0.266 | 0.011 |
| 37.000 | 0.242 | 0.021 | 46.000 | 0.272 | 0.009 |
| 38.000 | 0.237 | 0.024 | 47.000 | 0.275 | 0.008 |
| 39.000 | 0.251 | 0.016 | 48.000 | 0.281 | 0.007 |
| 40.000 | 0.255 | 0.015 | 49.000 | 0.283 | 0.007 |
| 41.000 | 0.257 | 0.014 | 50.000 | 0.281 | 0.007 |
| 42.000 | 0.269 | 0.010 | NaN | NaN | NaN |

TABLE S10: Correlation analysis between various global measures calculated at lag of 3 and the letter number sequencing test scores.

| Clustering | | | | | |
|---|---|---|---|---|---|
| **density** | **coefficient** | **p-value** | **density** | **coefficient** | **p-value** |
| 16.000 | -0.211 | 0.045 | 38.000 | -0.255 | 0.015 |
| 17.000 | -0.222 | 0.034 | 39.000 | -0.257 | 0.014 |
| 18.000 | -0.224 | 0.033 | 40.000 | -0.257 | 0.014 |
| 19.000 | -0.225 | 0.032 | 41.000 | -0.260 | 0.013 |
| 20.000 | -0.227 | 0.030 | 42.000 | -0.265 | 0.011 |
| 21.000 | -0.230 | 0.029 | 43.000 | -0.262 | 0.012 |
| 22.000 | -0.230 | 0.028 | 44.000 | -0.263 | 0.012 |
| 23.000 | -0.239 | 0.022 | 45.000 | -0.266 | 0.011 |
| 24.000 | -0.244 | 0.020 | 46.000 | -0.267 | 0.010 |
| 25.000 | -0.245 | 0.019 | 47.000 | -0.267 | 0.010 |
| 26.000 | -0.244 | 0.020 | 48.000 | -0.267 | 0.011 |
| 27.000 | -0.242 | 0.021 | 49.000 | -0.267 | 0.010 |
| 28.000 | -0.244 | 0.020 | 50.000 | -0.269 | 0.010 |
| 37.000 | -0.255 | 0.015 | NaN | NaN | NaN |
| **Transitivity** | | | | | |
| **density** | **coefficient** | **p-value** | **density** | **coefficient** | **p-value** |
| 25.000 | -0.224 | 0.033 | 38.000 | -0.254 | 0.015 |
| 26.000 | -0.225 | 0.032 | 39.000 | -0.256 | 0.014 |
| 27.000 | -0.229 | 0.029 | 40.000 | -0.257 | 0.014 |
| 28.000 | -0.233 | 0.026 | 41.000 | -0.259 | 0.013 |
| 29.000 | -0.235 | 0.025 | 42.000 | -0.265 | 0.011 |
| 30.000 | -0.238 | 0.023 | 43.000 | -0.262 | 0.012 |
| 31.000 | -0.240 | 0.022 | 44.000 | -0.263 | 0.012 |
| 32.000 | -0.242 | 0.021 | 45.000 | -0.266 | 0.011 |

Continued on next page

**TABLE S10 – continued from previous page**

| | | | | | |
|---|---|---|---|---|---|
| 33.000 | -0.243 | 0.020 | 46.000 | -0.267 | 0.011 |
| 34.000 | -0.248 | 0.018 | 47.000 | -0.268 | 0.010 |
| 35.000 | -0.249 | 0.017 | 48.000 | -0.268 | 0.010 |
| 36.000 | -0.250 | 0.017 | 49.000 | -0.268 | 0.010 |
| 37.000 | -0.254 | 0.015 | 50.000 | -0.270 | 0.010 |

TABLE S11: Correlation analysis between various global measures calculated at lag of 4 and the letter number sequencing test scores.

| **Global efficiency** | | | | | |
|---|---|---|---|---|---|
| **density** | **coefficient** | **p-value** | **density** | **coefficient** | **p-value** |
| 9.000 | -0.308 | 0.003 | 25.000 | -0.276 | 0.008 |
| 10.000 | -0.303 | 0.004 | 26.000 | -0.276 | 0.008 |
| 11.000 | -0.312 | 0.003 | 27.000 | -0.268 | 0.010 |
| 12.000 | -0.312 | 0.003 | 28.000 | -0.266 | 0.011 |
| 13.000 | -0.319 | 0.002 | 29.000 | -0.271 | 0.009 |
| 14.000 | -0.314 | 0.002 | 30.000 | -0.272 | 0.009 |
| 15.000 | -0.305 | 0.003 | 31.000 | -0.267 | 0.011 |
| 16.000 | -0.303 | 0.004 | 32.000 | -0.264 | 0.012 |
| 17.000 | -0.302 | 0.004 | 33.000 | -0.261 | 0.013 |
| 18.000 | -0.294 | 0.005 | 34.000 | -0.253 | 0.015 |
| 19.000 | -0.291 | 0.005 | 35.000 | -0.249 | 0.017 |
| 20.000 | -0.285 | 0.006 | 36.000 | -0.243 | 0.020 |
| 21.000 | -0.274 | 0.009 | 37.000 | -0.246 | 0.019 |
| 22.000 | -0.279 | 0.007 | 38.000 | -0.243 | 0.020 |
| 23.000 | -0.279 | 0.007 | 39.000 | -0.242 | 0.021 |
| 24.000 | -0.280 | 0.007 | NaN | NaN | NaN |
| **Local efficiency** | | | | | |
| **density** | **coefficient** | **p-value** | **density** | **coefficient** | **p-value** |
| 11.000 | -0.227 | 0.030 | 26.000 | -0.281 | 0.007 |
| 12.000 | -0.250 | 0.017 | 27.000 | -0.270 | 0.010 |
| 13.000 | -0.283 | 0.007 | 28.000 | -0.267 | 0.011 |
| 14.000 | -0.301 | 0.004 | 29.000 | -0.269 | 0.010 |
| 15.000 | -0.309 | 0.003 | 30.000 | -0.272 | 0.009 |
| 16.000 | -0.313 | 0.003 | 31.000 | -0.268 | 0.010 |

Continued on next page

**TABLE S11 – continued from previous page**

| | | | | | |
|---|---|---|---|---|---|
| 17.000 | -0.312 | 0.003 | 32.000 | -0.267 | 0.011 |
| 18.000 | -0.309 | 0.003 | 33.000 | -0.264 | 0.012 |
| 19.000 | -0.304 | 0.003 | 34.000 | -0.256 | 0.014 |
| 20.000 | -0.303 | 0.003 | 35.000 | -0.253 | 0.015 |
| 21.000 | -0.292 | 0.005 | 36.000 | -0.249 | 0.017 |
| 22.000 | -0.298 | 0.004 | 37.000 | -0.253 | 0.016 |
| 23.000 | -0.296 | 0.004 | 38.000 | -0.249 | 0.017 |
| 24.000 | -0.292 | 0.005 | 39.000 | -0.246 | 0.019 |
| 25.000 | -0.286 | 0.006 | 40.000 | -0.251 | 0.017 |

**Clustering**

| density | coefficient | p-value | density | coefficient | p-value |
|---|---|---|---|---|---|
| 11.000 | -0.213 | 0.043 | 31.000 | -0.274 | 0.008 |
| 12.000 | -0.224 | 0.033 | 32.000 | -0.275 | 0.008 |
| 13.000 | -0.236 | 0.024 | 33.000 | -0.277 | 0.008 |
| 14.000 | -0.242 | 0.021 | 34.000 | -0.276 | 0.008 |
| 15.000 | -0.251 | 0.016 | 35.000 | -0.274 | 0.009 |
| 16.000 | -0.254 | 0.015 | 36.000 | -0.275 | 0.008 |
| 17.000 | -0.263 | 0.012 | 37.000 | -0.276 | 0.008 |
| 18.000 | -0.257 | 0.014 | 38.000 | -0.275 | 0.008 |
| 19.000 | -0.260 | 0.013 | 39.000 | -0.279 | 0.007 |
| 20.000 | -0.257 | 0.014 | 40.000 | -0.280 | 0.007 |
| 21.000 | -0.255 | 0.015 | 41.000 | -0.281 | 0.007 |
| 22.000 | -0.265 | 0.011 | 42.000 | -0.280 | 0.007 |
| 23.000 | -0.269 | 0.010 | 43.000 | -0.281 | 0.007 |
| 24.000 | -0.270 | 0.010 | 44.000 | -0.281 | 0.007 |
| 25.000 | -0.267 | 0.010 | 45.000 | -0.279 | 0.007 |
| 26.000 | -0.272 | 0.009 | 46.000 | -0.280 | 0.007 |

Continued on next page

| | | | | | |
|---|---|---|---|---|---|
| 27.000 | -0.269 | 0.010 | 47.000 | -0.280 | 0.007 |
| 28.000 | -0.268 | 0.010 | 48.000 | -0.281 | 0.007 |
| 29.000 | -0.273 | 0.009 | 49.000 | -0.280 | 0.007 |
| 30.000 | -0.274 | 0.009 | 50.000 | -0.282 | 0.007 |
| **Transitivity** | | | | | |
| **density** | **coefficient** | **p-value** | **density** | **coefficient** | **p-value** |
| 23.000 | -0.254 | 0.015 | 37.000 | -0.278 | 0.008 |
| 24.000 | -0.257 | 0.014 | 38.000 | -0.277 | 0.008 |
| 25.000 | -0.257 | 0.014 | 39.000 | -0.281 | 0.007 |
| 26.000 | -0.263 | 0.012 | 40.000 | -0.281 | 0.007 |
| 27.000 | -0.263 | 0.012 | 41.000 | -0.282 | 0.007 |
| 28.000 | -0.263 | 0.012 | 42.000 | -0.281 | 0.007 |
| 29.000 | -0.268 | 0.010 | 43.000 | -0.283 | 0.007 |
| 30.000 | -0.269 | 0.010 | 44.000 | -0.284 | 0.006 |
| 31.000 | -0.271 | 0.009 | 45.000 | -0.282 | 0.007 |
| 32.000 | -0.271 | 0.009 | 46.000 | -0.283 | 0.007 |
| 33.000 | -0.274 | 0.009 | 47.000 | -0.283 | 0.007 |
| 34.000 | -0.275 | 0.008 | 48.000 | -0.284 | 0.006 |
| 35.000 | -0.274 | 0.008 | 49.000 | -0.283 | 0.007 |
| 36.000 | -0.277 | 0.008 | 50.000 | -0.285 | 0.006 |
| **Modularity** | | | | | |
| **density** | **coefficient** | **p-value** | **density** | **coefficient** | **p-value** |
| 32.000 | 0.227 | 0.030 | 42.000 | 0.280 | 0.007 |
| 33.000 | 0.233 | 0.026 | 43.000 | 0.285 | 0.006 |
| 34.000 | 0.256 | 0.014 | 44.000 | 0.290 | 0.005 |
| 35.000 | 0.252 | 0.016 | 45.000 | 0.290 | 0.005 |
| 36.000 | 0.269 | 0.010 | 46.000 | 0.287 | 0.006 |

Continued on next page

**TABLE S11 – continued from previous page**

| | | | | | |
|---|---|---|---|---|---|
| 37.000 | 0.275 | 0.008 | 47.000 | 0.286 | 0.006 |
| 38.000 | 0.275 | 0.008 | 48.000 | 0.288 | 0.006 |
| 39.000 | 0.288 | 0.006 | 49.000 | 0.287 | 0.006 |
| 40.000 | 0.281 | 0.007 | 50.000 | 0.283 | 0.007 |
| 41.000 | 0.284 | 0.006 | NaN | NaN | NaN |

TABLE S12: Correlation analysis between various global measures calculated at lag of 5 and the letter number sequencing test scores.

| Global efficiency | | | | | |
|---|---|---|---|---|---|
| **density** | **coefficient** | **p-value** | **density** | **coefficient** | **p-value** |
| 3.000 | -0.214 | 0.042 | 12.000 | -0.268 | 0.010 |
| 4.000 | -0.211 | 0.045 | 13.000 | -0.262 | 0.012 |
| 5.000 | -0.235 | 0.025 | 14.000 | -0.262 | 0.012 |
| 6.000 | -0.247 | 0.018 | 15.000 | -0.242 | 0.021 |
| 7.000 | -0.281 | 0.007 | 16.000 | -0.246 | 0.019 |
| 8.000 | -0.275 | 0.008 | 17.000 | -0.257 | 0.014 |
| 9.000 | -0.276 | 0.008 | 18.000 | -0.248 | 0.018 |
| 10.000 | -0.276 | 0.008 | 19.000 | -0.243 | 0.020 |
| 11.000 | -0.273 | 0.009 | 20.000 | -0.235 | 0.025 |
| **Local efficiency** | | | | | |
| **density** | **coefficient** | **p-value** | **density** | **coefficient** | **p-value** |
| 14.000 | -0.236 | 0.024 | 21.000 | -0.259 | 0.013 |
| 15.000 | -0.239 | 0.022 | 22.000 | -0.258 | 0.014 |
| 16.000 | -0.247 | 0.018 | 23.000 | -0.249 | 0.017 |
| 17.000 | -0.259 | 0.013 | 24.000 | -0.249 | 0.017 |
| 18.000 | -0.263 | 0.012 | 25.000 | -0.248 | 0.018 |
| 19.000 | -0.265 | 0.011 | 26.000 | -0.245 | 0.019 |
| 20.000 | -0.260 | 0.013 | 27.000 | -0.242 | 0.021 |
| **Clustering** | | | | | |
| **density** | **coefficient** | **p-value** | **density** | **coefficient** | **p-value** |
| 17.000 | -0.214 | 0.041 | 34.000 | -0.256 | 0.014 |
| 18.000 | -0.225 | 0.032 | 35.000 | -0.256 | 0.014 |
| 19.000 | -0.222 | 0.034 | 36.000 | -0.257 | 0.014 |
| 20.000 | -0.227 | 0.030 | 37.000 | -0.255 | 0.015 |

Continued on next page

**TABLE S12 – continued from previous page**

| | | | | | |
|---|---|---|---|---|---|
| 21.000 | -0.231 | 0.028 | 38.000 | -0.259 | 0.013 |
| 22.000 | -0.234 | 0.026 | 39.000 | -0.260 | 0.013 |
| 23.000 | -0.230 | 0.028 | 40.000 | -0.260 | 0.013 |
| 24.000 | -0.236 | 0.024 | 41.000 | -0.260 | 0.013 |
| 25.000 | -0.245 | 0.019 | 42.000 | -0.258 | 0.013 |
| 26.000 | -0.245 | 0.019 | 43.000 | -0.260 | 0.013 |
| 27.000 | -0.247 | 0.018 | 44.000 | -0.257 | 0.014 |
| 28.000 | -0.246 | 0.019 | 45.000 | -0.259 | 0.013 |
| 29.000 | -0.250 | 0.017 | 46.000 | -0.260 | 0.013 |
| 30.000 | -0.254 | 0.015 | 47.000 | -0.260 | 0.013 |
| 31.000 | -0.256 | 0.014 | 48.000 | -0.260 | 0.013 |
| 32.000 | -0.256 | 0.014 | 49.000 | -0.260 | 0.013 |
| 33.000 | -0.255 | 0.015 | 50.000 | -0.260 | 0.013 |

**Transitivity**

| **density** | **coefficient** | **p-value** | **density** | **coefficient** | **p-value** |
|---|---|---|---|---|---|
| 23.000 | -0.216 | 0.039 | 37.000 | -0.250 | 0.017 |
| 24.000 | -0.221 | 0.035 | 38.000 | -0.254 | 0.015 |
| 25.000 | -0.229 | 0.029 | 39.000 | -0.255 | 0.015 |
| 26.000 | -0.230 | 0.028 | 40.000 | -0.256 | 0.014 |
| 27.000 | -0.233 | 0.026 | 41.000 | -0.257 | 0.014 |
| 28.000 | -0.233 | 0.026 | 42.000 | -0.256 | 0.014 |
| 29.000 | -0.238 | 0.023 | 43.000 | -0.258 | 0.014 |
| 30.000 | -0.241 | 0.021 | 44.000 | -0.257 | 0.014 |
| 31.000 | -0.245 | 0.019 | 45.000 | -0.258 | 0.014 |
| 32.000 | -0.246 | 0.019 | 46.000 | -0.260 | 0.013 |
| 33.000 | -0.247 | 0.018 | 47.000 | -0.260 | 0.013 |
| 34.000 | -0.248 | 0.018 | 48.000 | -0.261 | 0.012 |

Continued on next page

**TABLE S12 – continued from previous page**

| | | | | | |
|---|---|---|---|---|---|
| 35.000 | -0.249 | 0.017 | 49.000 | -0.261 | 0.013 |
| 36.000 | -0.250 | 0.017 | 50.000 | -0.260 | 0.013 |

**Modularity**

| density | coefficient | p-value | density | coefficient | p-value |
|---|---|---|---|---|---|
| 39.000 | 0.248 | 0.018 | 45.000 | 0.262 | 0.012 |
| 40.000 | 0.249 | 0.017 | 46.000 | 0.264 | 0.011 |
| 41.000 | 0.259 | 0.013 | 47.000 | 0.265 | 0.011 |
| 42.000 | 0.267 | 0.011 | 48.000 | 0.264 | 0.011 |
| 43.000 | 0.264 | 0.012 | 49.000 | 0.261 | 0.013 |
| 44.000 | 0.264 | 0.011 | 50.000 | 0.261 | 0.012 |

TABLE S13: Correlation analysis between various global measures calculated at lag of 6 and the letter number sequencing test scores.

| **Global efficiency** | | | | | |
|---|---|---|---|---|---|
| **density** | **coefficient** | **p-value** | **density** | **coefficient** | **p-value** |
| 4.000 | -0.235 | 0.025 | 8.000 | -0.259 | 0.013 |
| 5.000 | -0.239 | 0.022 | 9.000 | -0.264 | 0.012 |
| 6.000 | -0.242 | 0.021 | 10.000 | -0.261 | 0.012 |
| 7.000 | -0.253 | 0.016 | NaN | NaN | NaN |
| **Modularity** | | | | | |
| **density** | **coefficient** | **p-value** | **density** | **coefficient** | **p-value** |
| 25.000 | 0.235 | 0.025 | 35.000 | 0.259 | 0.013 |
| 26.000 | 0.304 | 0.003 | 36.000 | 0.275 | 0.008 |
| 27.000 | 0.242 | 0.021 | 37.000 | 0.268 | 0.010 |
| 28.000 | 0.234 | 0.026 | 38.000 | 0.258 | 0.014 |
| 29.000 | 0.253 | 0.015 | 39.000 | 0.248 | 0.018 |
| 30.000 | 0.334 | 0.001 | 40.000 | 0.246 | 0.019 |
| 31.000 | 0.288 | 0.006 | 41.000 | 0.234 | 0.025 |
| 32.000 | 0.299 | 0.004 | 42.000 | 0.231 | 0.028 |
| 33.000 | 0.277 | 0.008 | 43.000 | 0.227 | 0.031 |
| 34.000 | 0.274 | 0.009 | NaN | NaN | NaN |

TABLE S14: Correlation analysis between various global measures calculated at lag of 7 and the letter number sequencing test scores.

| **Modularity** | | | | | |
|---|---|---|---|---|---|
| **density** | **coefficient** | **p-value** | **density** | **coefficient** | **p-value** |
| 24.000 | 0.312 | 0.003 | 29.000 | 0.305 | 0.003 |
| 25.000 | 0.287 | 0.006 | 30.000 | 0.297 | 0.004 |
| 26.000 | 0.332 | 0.001 | 31.000 | 0.274 | 0.009 |
| 27.000 | 0.312 | 0.003 | 32.000 | 0.274 | 0.009 |
| 28.000 | 0.269 | 0.010 | 33.000 | 0.254 | 0.015 |

TABLE S15: Correlation analysis between various global measures calculated at lag of 1 and the symbol digit modalities test scores.

| **Clustering** | | | | | |
|---|---|---|---|---|---|
| **density** | **coefficient** | **p-value** | **density** | **coefficient** | **p-value** |
| 28.000 | -0.222 | 0.034 | 40.000 | -0.241 | 0.021 |
| 29.000 | -0.225 | 0.032 | 41.000 | -0.236 | 0.024 |
| 30.000 | -0.226 | 0.031 | 42.000 | -0.237 | 0.024 |
| 31.000 | -0.230 | 0.028 | 43.000 | -0.238 | 0.023 |
| 32.000 | -0.228 | 0.029 | 44.000 | -0.238 | 0.023 |
| 33.000 | -0.229 | 0.029 | 45.000 | -0.239 | 0.023 |
| 34.000 | -0.229 | 0.029 | 46.000 | -0.239 | 0.022 |
| 35.000 | -0.229 | 0.029 | 47.000 | -0.240 | 0.022 |
| 36.000 | -0.231 | 0.027 | 48.000 | -0.243 | 0.020 |
| 37.000 | -0.233 | 0.026 | 49.000 | -0.242 | 0.021 |
| 38.000 | -0.235 | 0.025 | 50.000 | -0.244 | 0.020 |
| 39.000 | -0.238 | 0.023 | NaN | NaN | NaN |
| **Transitivity** | | | | | |
| **density** | **coefficient** | **p-value** | **density** | **coefficient** | **p-value** |
| 25.000 | -0.215 | 0.041 | 38.000 | -0.241 | 0.021 |
| 26.000 | -0.216 | 0.040 | 39.000 | -0.245 | 0.019 |
| 27.000 | -0.219 | 0.037 | 40.000 | -0.247 | 0.018 |
| 28.000 | -0.223 | 0.034 | 41.000 | -0.242 | 0.021 |
| 29.000 | -0.228 | 0.030 | 42.000 | -0.242 | 0.021 |
| 30.000 | -0.228 | 0.030 | 43.000 | -0.242 | 0.021 |
| 31.000 | -0.232 | 0.027 | 44.000 | -0.243 | 0.020 |
| 32.000 | -0.232 | 0.027 | 45.000 | -0.244 | 0.020 |
| 33.000 | -0.234 | 0.026 | 46.000 | -0.245 | 0.019 |
| 34.000 | -0.235 | 0.025 | 47.000 | -0.245 | 0.019 |

Continued on next page

**TABLE S15 – continued from previous page**

| | | | | | |
|---|---|---|---|---|---|
| 35.000 | -0.236 | 0.025 | 48.000 | -0.248 | 0.018 |
| 36.000 | -0.239 | 0.023 | 49.000 | -0.248 | 0.018 |
| 37.000 | -0.240 | 0.022 | 50.000 | -0.250 | 0.017 |

TABLE S16: Correlation analysis between various global measures calculated at lag of 5 and the Hopkins verbal learning test-revised test scores.

| **Global efficiency** | | | | | |
| --- | --- | --- | --- | --- | --- |
| **density** | **coefficient** | **p-value** | **density** | **coefficient** | **p-value** |
| 7.000 | -0.221 | 0.035 | 14.000 | -0.234 | 0.025 |
| 8.000 | -0.224 | 0.033 | 15.000 | -0.225 | 0.032 |
| 9.000 | -0.217 | 0.038 | 16.000 | -0.227 | 0.030 |
| 10.000 | -0.224 | 0.033 | 17.000 | -0.217 | 0.039 |
| 11.000 | -0.227 | 0.031 | 18.000 | -0.211 | 0.045 |
| 12.000 | -0.232 | 0.027 | 19.000 | -0.210 | 0.045 |
| 13.000 | -0.237 | 0.024 | NaN | NaN | NaN |

TABLE S17: Correlation analysis between various global measures calculated at lag of 7 and the Benton's judgment of line orientation test scores.

| **Global efficiency** | | | | | |
|---|---|---|---|---|---|
| **density** | **coefficient** | **p-value** | **density** | **coefficient** | **p-value** |
| 5.000 | 0.221 | 0.035 | 9.000 | 0.236 | 0.025 |
| 6.000 | 0.219 | 0.037 | 10.000 | 0.235 | 0.025 |
| 7.000 | 0.227 | 0.031 | 11.000 | 0.229 | 0.029 |
| 8.000 | 0.226 | 0.031 | NaN | NaN | NaN |
| **Local efficiency** | | | | | |
| **density** | **coefficient** | **p-value** | **density** | **coefficient** | **p-value** |
| 8.000 | 0.232 | 0.027 | 13.000 | 0.257 | 0.014 |
| 9.000 | 0.236 | 0.025 | 14.000 | 0.256 | 0.015 |
| 10.000 | 0.243 | 0.020 | 15.000 | 0.260 | 0.013 |
| 11.000 | 0.242 | 0.021 | 16.000 | 0.269 | 0.010 |
| 12.000 | 0.244 | 0.020 | 17.000 | 0.276 | 0.008 |
| **Clustering** | | | | | |
| **density** | **coefficient** | **p-value** | **density** | **coefficient** | **p-value** |
| 7.000 | 0.230 | 0.028 | 29.000 | 0.236 | 0.024 |
| 8.000 | 0.220 | 0.036 | 30.000 | 0.239 | 0.023 |
| 9.000 | 0.212 | 0.044 | 31.000 | 0.239 | 0.023 |
| 10.000 | 0.227 | 0.031 | 32.000 | 0.243 | 0.020 |
| 11.000 | 0.222 | 0.035 | 33.000 | 0.238 | 0.023 |
| 12.000 | 0.226 | 0.032 | 34.000 | 0.237 | 0.024 |
| 13.000 | 0.228 | 0.030 | 35.000 | 0.241 | 0.022 |
| 14.000 | 0.228 | 0.029 | 36.000 | 0.242 | 0.021 |
| 15.000 | 0.233 | 0.026 | 37.000 | 0.246 | 0.019 |
| 16.000 | 0.232 | 0.027 | 38.000 | 0.245 | 0.019 |
| 17.000 | 0.235 | 0.025 | 39.000 | 0.243 | 0.020 |

Continued on next page

**TABLE S17 – continued from previous page**

| | | | | | |
|---|---|---|---|---|---|
| 18.000 | 0.234 | 0.026 | 40.000 | 0.245 | 0.019 |
| 19.000 | 0.242 | 0.021 | 41.000 | 0.247 | 0.018 |
| 20.000 | 0.241 | 0.021 | 42.000 | 0.247 | 0.018 |
| 21.000 | 0.243 | 0.020 | 43.000 | 0.248 | 0.018 |
| 22.000 | 0.246 | 0.019 | 44.000 | 0.249 | 0.018 |
| 23.000 | 0.242 | 0.021 | 45.000 | 0.248 | 0.018 |
| 24.000 | 0.241 | 0.021 | 46.000 | 0.247 | 0.019 |
| 25.000 | 0.234 | 0.025 | 47.000 | 0.247 | 0.018 |
| 26.000 | 0.239 | 0.023 | 48.000 | 0.249 | 0.017 |
| 27.000 | 0.239 | 0.023 | 49.000 | 0.250 | 0.017 |
| 28.000 | 0.234 | 0.025 | 50.000 | 0.252 | 0.016 |
| **Transitivity** | | | | | |
| **density** | **coefficient** | **p-value** | **density** | **coefficient** | **p-value** |
| 13.000 | 0.219 | 0.037 | 32.000 | 0.246 | 0.019 |
| 14.000 | 0.224 | 0.032 | 33.000 | 0.243 | 0.020 |
| 15.000 | 0.227 | 0.030 | 34.000 | 0.243 | 0.020 |
| 16.000 | 0.231 | 0.027 | 35.000 | 0.246 | 0.019 |
| 17.000 | 0.234 | 0.025 | 36.000 | 0.245 | 0.019 |
| 18.000 | 0.236 | 0.024 | 37.000 | 0.248 | 0.018 |
| 19.000 | 0.241 | 0.021 | 38.000 | 0.248 | 0.018 |
| 20.000 | 0.243 | 0.020 | 39.000 | 0.246 | 0.019 |
| 21.000 | 0.244 | 0.020 | 40.000 | 0.248 | 0.018 |
| 22.000 | 0.248 | 0.018 | 41.000 | 0.249 | 0.017 |
| 23.000 | 0.245 | 0.019 | 42.000 | 0.249 | 0.017 |
| 24.000 | 0.244 | 0.020 | 43.000 | 0.249 | 0.017 |
| 25.000 | 0.242 | 0.021 | 44.000 | 0.249 | 0.017 |
| 26.000 | 0.245 | 0.019 | 45.000 | 0.249 | 0.018 |

Continued on next page

**TABLE S17 – continued from previous page**

| | | | | | |
|---|---|---|---|---|---|
| 27.000 | 0.245 | 0.019 | 46.000 | 0.248 | 0.018 |
| 28.000 | 0.243 | 0.020 | 47.000 | 0.248 | 0.018 |
| 29.000 | 0.244 | 0.020 | 48.000 | 0.250 | 0.017 |
| 30.000 | 0.244 | 0.020 | 49.000 | 0.250 | 0.017 |
| 31.000 | 0.244 | 0.020 | 50.000 | 0.251 | 0.016 |

#### III. PARTICIPANTS - DIVISION INTO SUBGROUPS BASED ON MEDICATION STATUS.

| | **MED (n = 64)** | **non-MED (n = 31)** | **MED vs non-MED (p value)** |
|---|---|---|---|
| **Age (years)** | 68.7 (10.4) | 66.6 (10.6) | 0.38 |
| **Gender (% male)** | 76.6% | 51.6% | 0.02 |
| **Education (years)** | 14.7 (2.8) | 16.5 (2.6) | 0.01 |
| **UPDRS-III test scores** | 22.0 (11.3) | 19.7 (9.2) | 0.33 |
| **HY stage (1-2)** | 16 - 48 | 11 - 20 | – |
| **LEDD (dose)** | 405.3 (207.0) | – | – |
| **Cognitive status (% MCI)** | 20.3% | 19.4% | – |

TABLE S18. **Characteristics of the sample - division into subgroups based on the medication status**. Means are followed by standard deviation in parentesis. Permutation tests with 10000 permutations were used to compare groups for age, gender, education and UPDRS-III scores. MED, Parkinson's disease patients on levodopa medication; non-MED, Parkinson's disease patients off levodopa medication; UPDRS-III, Unified Parkinson's disease rating scale–Part III; HY stage, Hoehn and Yahr stage; LEDD, levodopa equivalent dose; MCI, mild cognitive impairment.

#### IV. PARTICIPANTS - DIVISION INTO SUBGROUPS BASED ON COGNITIVE STATUS.

| | **PDCN (n = 76)** | **PDMCI (n = 19)** | **PDCN vs PDMCI (p value)** |
|---|---|---|---|
| **Age (years)** | 67.3 (10.6) | 70.9 (9.4) | 0.18 |
| **Gender (% male)** | 64.5% | 84.2% | 0.11 |
| **Education (years)** | 15.4 (2.9) | 14.8 (2.9) | 0.48 |
| **UPDRS-III test scores** | 20.7 (10.5) | 23.7 (11.2) | 0.28 |
| **HY stage (1-2)** | 22 - 54 | 5 - 14 | – |
| **LEDD (dose)** | 396.4 (199.2) | 440.3 (240.9) | – |
| **LEDD (% medicated)** | 67.1% | 68.4% | – |

TABLE S19. **Characteristics of the sample - division into subgroups based on cognitive status**. Means are followed by standard deviation in parentesis. Permutation tests with 10000 permutations were used to compare groups for age, gender, education and UPDRS-III scores. PDCN, Parkinson's disease cognitively normal; PDMCI, Parkinson's disease with mild cognitive impairment; UPDRS-III, Unified Parkinson's disease rating scale–Part III; HY stage, Hoehn and Yahr stage; LEDD, levodopa equivalent dose.
